# The immunophilin protein FKBPL and its peptide derivatives are novel regulators of vascular integrity and inflammation via NF-κB signaling

**DOI:** 10.1101/2021.02.24.431422

**Authors:** Stephanie Annett, Shaun Spence, Carolina Garciarena, Ciaran Campbell, Margaret Dennehy, Clive Drakeford, Jacqueline Lai, Jennifer Dowling, Gillian Moore, Anita Yakkundi, Amy Short, Danny Sharpe, Fiona Furlong, James S. O’Donnell, Gianpiero Cavalleri, Steve Kerrigan, Irina G. Tikhonova, Pauline Johnson, Adrien Kissenpfennig, Tracy Robson

## Abstract

A breakdown in vascular integrity and excessive inflammation are hallmarks of serious pathological conditions including sepsis, acute respiratory distress syndrome (ARDs) and most recently, severe COVID-19. FK506 – binding protein like (FKBPL) is a member of the immunophilin protein superfamily with potent anti-tumor activity through inhibition of angiogenesis and cancer stemness. An FKBPL-based 23mer peptide, ALM201, displayed a good safety and pharmacokinetic profile in a Phase 1a oncology clinical trial and was subsequently designated orphan drug status by the FDA in ovarian cancer. Here we describe a novel role for FKBPL and its peptides in regulating vascular integrity and cytokine production though modulating NF-κB signaling. FKBPL knockdown promoted endothelial cell barrier permeability, which was further exacerbated upon stimulation with lipopolysaccharide (LPS) and accompanied by increased expression of TNF mRNA and phosphorylation of p65(RelA). Whilst treatment with the FKBPL based pre-clinical peptide, AD-01, increased VE-cadherin endothelial tight junctions following LPS stimulation. Bone marrow derived macrophages (BMDM) from FKBPL haploinsufficient mice (*Fkbpl*^+/−^) also demonstrated increased phosphorylation of p65(RelA) in response to LPS stimulation compared to wild-type mice. Furthermore, treatment with AD-01 inhibited p65(RelA) phosphorylation following LPS stimulation resulting in reduced NF-κB target gene expression and proinflammatory cytokine production. In an *in vivo* LPS survival model, *Fkbpl*^+/−^ mice have reduced survival compared to wild-type mice. Moreover, treatment of wild-type mice with the clinical FKBPL-based peptide, ALM201, following LPS injection resulted in a 100% survival rate in mice at experimental endpoint, as well as an abrogation of production of pro-inflammatory cytokines, TNF and IL-6, in peritoneal lavage washings. Analysis of human genetic biobanks found an association between common genetic variants associated with FKBPL and traits associated with inflammatory disorders such as psoriasis, rheumatoid arthritis and high lymphocyte count. In summary, for the first time, we describe a novel role for FKBPL as a regulator of inflammation and vascular integrity through modulating NF-κB signaling and FKBPL based therapies demonstrate potent anti-inflammatory activity.

## Introduction

Sepsis, acute respiratory distress syndrome (ARDS), influenza and, most recently, COVID-19 can progress to systemic inflammatory response syndrome (SIRS) (1, 2). SIRS develops when the initial, appropriate innate immune responses become amplified and then dysregulated (3). High levels of pathogen-associated molecular pattern (PAMP) mediators, such as lipopolysaccharide (LPS) or viral RNA, results in a large increase in systemic inflammatory mediators which then contributes to organ failure (2, 4). In addition, endothelial barrier integrity is disrupted in SIRS causing extravascular fluid leakage, disseminated intravascular coagulation and septic shock (2, 3). Enhanced vascular permeability and systemic inflammation are also the hallmarks of many chronic inflammatory conditions, such as psoriasis, inflammatory bowel disease and rheumatoid arthritis (5–7). We hypothesize that strategic targeting of both vascular permeability and systemic inflammation has potential benefits on survival and quality of life in both acute and chronic inflammatory syndromes.

The nuclear factor-κB (NF-κB) family of inducible transcription factors are master regulators of immune responses (8). Upon activation of the canonical signaling pathway, a multi-subunit IκB kinase (IKK) complex phosphorylates IκBα at two N-terminal serines and, thereby, triggers ubiquitin-dependent IκBα in the proteasome resulting in rapid and transient nuclear translocation of canonical NF-κB members, predominately p50/RelA(p65) dimers (8). This results in the expression of proinflammatory genes, the activation of the inflammasome and loss of vascular integrity in endothelial cells (8–11). Unselective pharmacological inhibition of NF-κB, across cell types, results in multiple adverse effects; a major hindrance in drug development (12). The nuclear translocation of NF-κB dimers is dependent upon a dynein/dynactin motor – immunophilin complex, indicating that immunophilins can fine tune the signaling cascade (13, 14). Therefore, non-direct modulation of NF-κB signaling through modulating immunophilins may be a less toxic therapeutic target.

Immunophilins are a highly conserved intracellular protein superfamily best known for their role in binding the immunosuppressive drugs, FK506 (tacrolimus), rapamycin (sirolimus) and cyclosporine A (CsA) and they consist of three main groups; cyclophilins, FK506-binding proteins (FKBPs) and parvulins (15). A key feature of the family is the presence of functional and non-functional peptidyl prolyl isomerases (PPIase) domains which have cis-trans isomerization activity at the X-Pro peptide bond (16). The members of the FKBP family containing tetratricopeptide repeat (TPR) domains, FKBP51 and FKB52, exhibit opposing inhibitory and stimulatory activity on NF-κB signaling through interactions with the p65 complex (17). The immunophilin, FK506-binding protein like (FKBPL), is a divergent member of the FKBP family and was first identified by screening for genes involved in the radiation response (18, 19). FKBPL also contains three TPR domains in the C terminal region, important for interactions with heat shock protein 90 (Hsp90), and a non-functional PPIase domain at the N terminal (20). In a complex with Hsp90, FKBPL stabilizes the cyclin-dependent kinase inhibitor, p21, and complexes with the estrogen receptor (ER), androgen receptor and glucocorticoid receptor (21–24). In addition, FKBPL is a secreted anti-angiogenic protein and the cell surface receptor, CD44, is a potential target for its activity by binding to the N terminal region of FKBPL (25, 26). In support of a role for FKBPL in angiogenesis, FKBPL knockout mice are embryonically lethal and FKBPL haploinsufficient (*Fkbpl^+/−^*) embryos display vascular irregularities, suggesting a critical role for FKBPL in developmental angiogenesis (27). In the cancer setting, FKBPL also demonstrated dual anti-angiogenic and anti-cancer stem cell (CSC) activity (28–31). Indeed, high intra-tumor FKBPL levels are a marker of improved prognosis in breast and ovarian cancer (32, 33). *In vitro* and *in vivo* knockdown of FKBPL expands the CSC subpopulation by increasing expression of the pluripotency transcription factors (*NANOG, OCT4, SOX2*) (29). Notably, an increase in *NFKBP1*, which encodes for the p105 NF-κB pre cursor subunit, was also observed when FKBPL was knocked down in ovarian cancer (33). The highly potent anti-angiogenic and anti-CSC activity of FKBPL is due to a unique sequence within the N-terminal region, independent of the TPR domains (34). A 24-residue peptide, AD-01, comprising amino acids 34–58 of FKBPL was developed demonstrating potent anti-tumor activity and a more stable 23-residue peptide, ALM201, was selected as the clinical drug candidate. ALM201 elicited equipotent activity to AD-01 (33, 35). ALM201 lacked cytotoxicity and displayed an excellent safety profile in a Phase 1a, first-in-man, dose-escalation clinical trial in patients with ovarian cancer and other solid tumors (EudraCT number: 2014-001175-31) and was subsequently designated orphan drug status by the FDA in ovarian cancer (36).

Here for the first time, we discover a novel role for FKBPL and its peptide derivatives in the regulation of inflammation via modulation of NF-κB signaling. In macrophages, FKBPL regulates secretion of pro-inflammatory cytokines and in endothelial cells, FKBPL has a role in maintaining vascular integrity in response to inflammation. Moreover, FKBPL based peptides, previously shown to be safe in humans, have potent anti-inflammatory activity and abrogate LPS lethality in *in vivo* models. Finally, we demonstrate that genetic variation in human FKBPL is associated with chronic inflammatory disorders such as psoriasis and rheumatoid arthritis, supporting a protective role for this protein in pathological conditions associated with inflammation.

## Materials and Methods

### Cell Culture

Human microvascular endothelial cells (HMEC-1, ATTC) were cultured at 37°C in a humidified atmosphere of 95% O2/5% CO_2_. Culture media contained MCDB-131 (Invitrogen, Ireland), 10 ng/ml recombinant human EGF (Roche, Ireland), 1 μg/ml hydrocortisone (Sigma, Ireland), 10 nM of L-glutamine (Invitrogen, Ireland), 10% Fetal calf serum (FCS) (Invitrogen, Ireland), and 1% penicillin/streptomycin (Invitrogen, Ireland). THP-1 cells were a kind gift from Prof James O’Donnell and were cultured in RPMI media (Invitrogen, Ireland) with 10% FCS and 1% penicillin/streptomycin at a temperature of 37°C, 95% air and 5% CO_2_. Suspension THP-1 cells were maintained at a density 3-10 x 10^5^ cells/ml in non-adherent plastic (Sarstedt, Germany), were cultured for two weeks prior to use and not maintained beyond passage 20. L929 cells were a kind gift from Dr. Jennifer Dowling and conditioned media was prepared to promote differentiation of bone marrow progenitors to bone marrow derived macrophages (BMDMs). Monthly testing ensured cells were Mycoplasma free. Cells were treated with LPS 0111:B4 (Sigma, Ireland) and/or PBS vehicle control (Invitrogen, Ireland), AD-01 (NH2-QIRQQPRDPPTETLELEVSPDPAS-OH) (Almac Group, United Kingdom), ALM201 (NH2-IRQQPRDPPTETLELEVSPDPAS-OH) (Almac group, United Kingdom) and recombinant FKBPL protein (ab98117; Abcam). All experiments were carried out at 37°C in a humidified atmosphere of 95% O_2_/5% CO_2._

### Transgenic mouse colony

The *Fkbpl^+/−^* (tm1.1(KOMP)Wtsi) mice were obtained from the trans-NIH Knock-Out Mouse Project (KOMP) Repository (www.komp.org) and were housed in the Biological Resource Unit at RCSI University of Medicine and Health Sciences (EPA license no. 693) and Queen’s University Belfast. The CD44^−/−^ mice (37) where housed and bred at the University of British Columbia as approved by the University Animal Care Committee in accordance with the Canadian Council of Animal Care guidelines for ethical animal research (A15-213, A19-159). All animal experiments were approved by the Health Products Research Authority or UK Home Office and RCSI Research and Ethics Committee under Project License PPL 1403 and Individual license numbers IAN I181 and IAN I193. Experiments were carried out in accordance with Directive 2010/63/EU.

### Bone marrow derived macrophages

Bone marrow was removed from the femur and tibia bone by inserting the top of a syringe into the orifice of the bone and flushing with PBS. The extracted bone marrow was resuspended in red blood cell lysis buffer (Sigma, Ireland) and incubated for 2 min at room temperature. PBS was added to stop the reaction and cells were resuspended in differentiation media (DMEM with 10% (v/v) FBS, 1% (v/v) penicillin and streptomycin and M-CSF (20% (v/v) L929 mouse fibroblast supernatant). Cells were then strained through a 70 μm filter and plated on petri dishes and incubated in 37°C in a humidified atmosphere of 95% O2/5% CO_2_ for 6 days, with media changed on day 3. Cells were then plated for experiments at a concentration of 5 x 10^5^ cells per ml in complete DMEM media on tissue culture plates for 24 h and treatments added as described.

### siFKBPL transfection

HMEC-1 cells were grown until 80% confluence in a 6 well plate and then transfected using 100 nM ON TARGETplus SMARTpool siFKBPL (Horizon Discovery, UK) or siNon-targeted negative control (Horizon Discovery, UK) and Lipofectamine (10 μl/well) (Invitrogen, UK) in Opti-MEM I Reduced Serum Medium (Invitrogen, Ireland). Cells were incubated at 37°C in a humidified atmosphere of 95% O_2_/5% CO_2_, for 8 h. Cells were seeded in Transwell support inserts X-CELLigence plates for *in vitro* endothelial barrier assays or cultured for a further 24 h and for gene expression analysis or 72 h for protein expression analysis in complete media.

### In vitro Evan’s blue permeability assay

Endothelial cell barrier permeability was determined as described previously (38), with minor modifications. Transfected HMEC-1 cells (2.5 x 10^5^ cells/ml in complete media) were seeded for 72 h on polycarbonate membrane Transwell permeable inserts (Costar, 3 μm pore size, 12 mm diameter, Sigma Ireland) and incubated for three days. Fresh growth media was added to the bottom chambers, and media was removed from the top chamber and replaced with 0.67 mg/ml Evans Blue (Sigma, Ireland) with 4% bovine serum albumin (BSA; Sigma, Ireland) for 10 min. A change in endothelial cell barrier permeability was determined by assessment of the increase in absorbance at 650 nm in the bottom chamber due to Evans Blue-BSA permeating the endothelial cell layer. Experiments were performed and permeability (percentage) was determined compared to untreated cells.

### In vivo Evan’s blue (Miles Assay)

A 0.5% sterile solution of Evans blue (Sigma, Ireland) was prepared in PBS and 200 μl slowly injected into the tail vein of mice. After 30 min the mice were sacrificed through cervical dislocation and organs collected and weighed. 500 μl formamide (Sigma, Ireland) was added to the organs and heated to 55°C in a heat block for 24 h. The formamide/Evan’s blue mixture was centrifuged to pellet any tissue and the supernatant measured at 610 nm, using formamide as a blank. The amount of Evan’s blue extravasated per mg of tissue was then calculated as outlined in (39).

### X-CELLigence

Using the X-CELLigence system (Roche, Welwyn Garden City, UK) to assess impedance, 3 × 10^4^ HMEC-1 cells were seeded into an E-16 multi-well plate (Roche) in triplicate and incubated for 72 h. LPS (Sigma, Ireland) and/or AD-01 (100 nM) was added to the relevant wells. Impedance measurements were taken at 10 min intervals and cell index normalized to the point immediately prior to addition of LPS (1 μg/ml and 10 μg/ml) and PBS or AD-01 (1 nM and 100 nM).

### Immunofluorescence

3 x 10^5^ HMEC-1 cells/well were seeded in a 6-well plate with inserted glass cover slips fixed using nail varnish. Cells were incubated for 24 h and then sheared at 10 dyn/cm^2^ for a further 24 h. LPS (10 μg/ml) and AD-01 (100 nM) was added to the cells under shear conditions for 2 h. The cells were fixed for 10 min using 4% paraformaldehyde at room temperature and washed twice with PBS for 10 min. Triton 0.01% in PBS was added for 20 min and cells washed twice with PBS for 10 min. A blocking solution of 1% BSA in PBS was added for 1 h at room temperature. Anti VE cadherin was added (sc-9989, Santa Cruz, USA) in a 1:50 dilution in blocking buffer for 1 h at room temperature. Cells were washed with PBS twice for 10 min. A 1:1000 dilution of anti-mouse Alexa 488 (Invitrogen, Ireland) was added for 45 min at room temperature. Cells were washed twice for 10 min with PBS and a cover slip was mounted to a glass slide with a mounting solution containing DAPI (Abcam). Cells were visualized using confocal microscopy and VE cadherin surface expression quantified using ImageJ 1x (Image J, NIH)

### Inflammasome activation

Differentiated BMDM were treated with LPS (100 ng/ml) and incubated for 3 h. The media was then removed and PBS or AD-01 (1 nM) was then added for 1 h in serum free media. To induce IL-1β cleavage, adenosine triphosphate (ATP) (5 μM, Invivogen, USA) was added for 45 min. Supernatants were collected and cells were lysed in RIPA buffer and analyzed by Western blot or ELISA, as described.

### Real-time PCR

Cells were washed with PBS followed by lysis and homogenization with 1 ml of TRIZOL (Sigma-Aldrich, Ireland) per 10 cm^2^ culture and RNA extracted as per manufacturer’s protocol. RNA concentration and purity were determined by 260λ/280λ and 260λ/280λ ratios respectively. cDNA was synthesized from 100 ng-200 ng of RNA using M-MLV reverse transcriptase (Invitrogen, Ireland) in a thermocyler. Gene expression was determined by quantitative reverse transcription PCR (qPCR). Gene specific primers (Table 1; IDT, USA) were diluted 1/10 from stock 100 μM concentrations and PCR reaction was completed using a 7500 real time PCR system (ThermoFisher, USA) and Sybr Green (Invitrogen, Ireland). Changes in expression of genes of interest from untreated cells, or treatments were quantified by fold change-ΔΔCT from housekeeping genes β-Actin and GADPH (2–ΔΔCt).

**Table 1.**
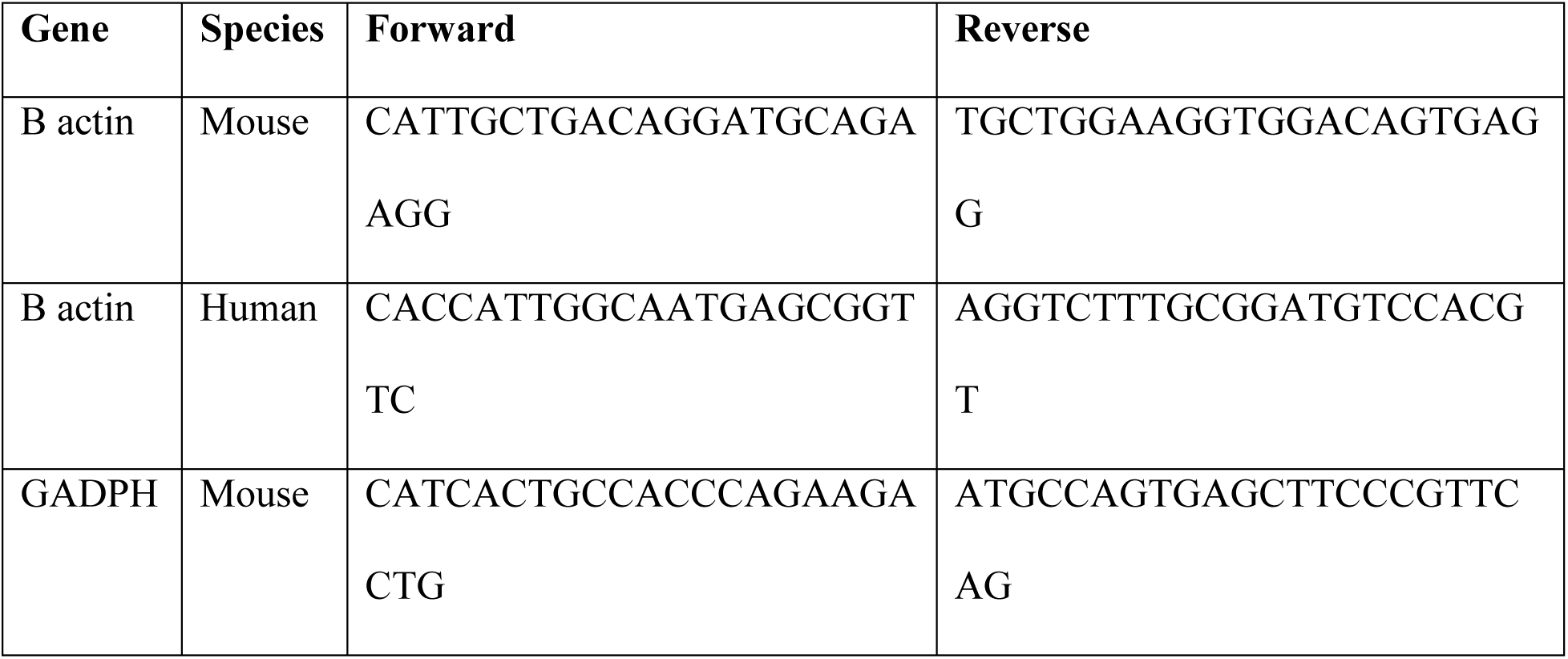
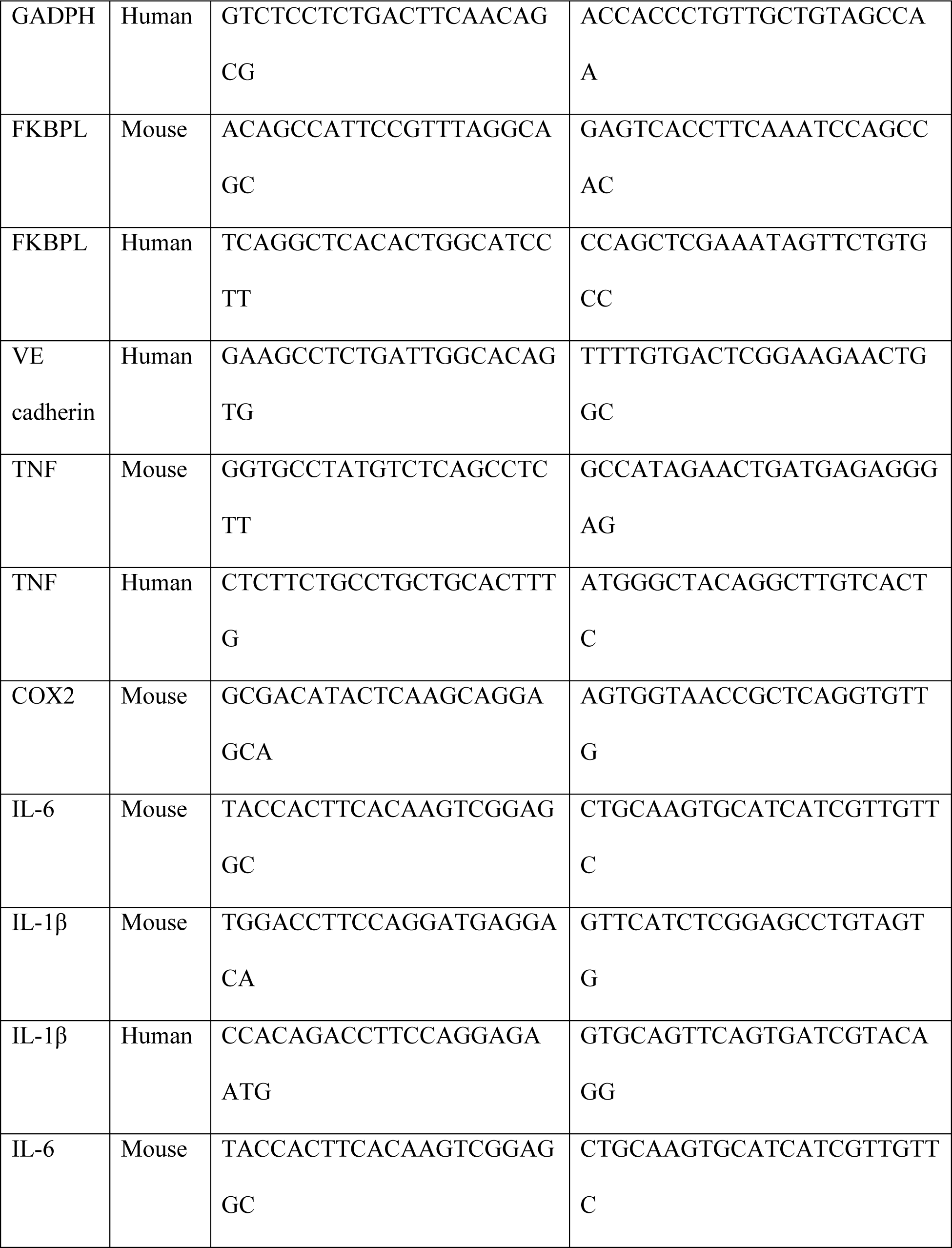
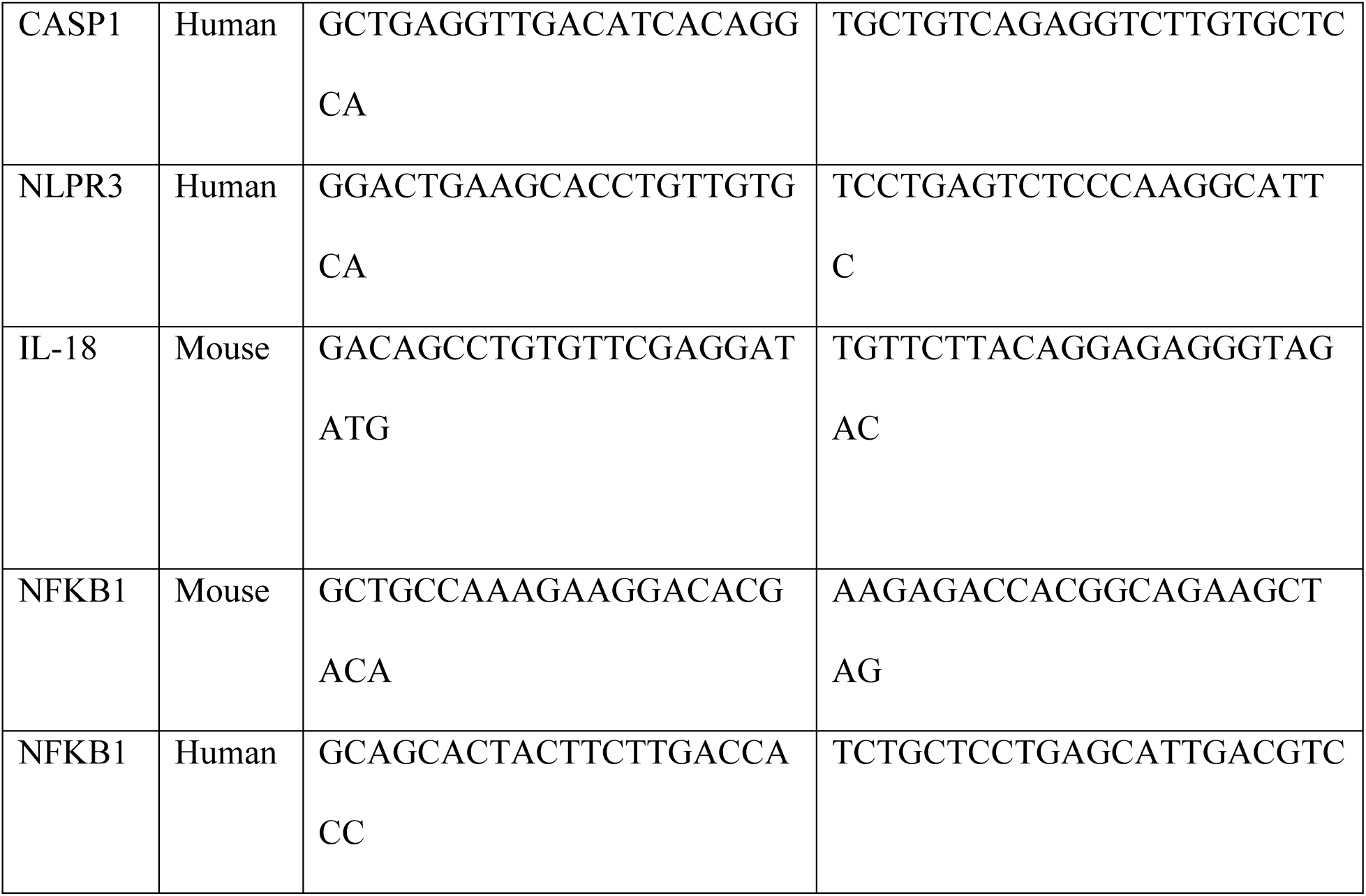
Primer sequences used for gene expression analysis

### Western blot

Cells were lysed on ice with RIPA buffer (Sigma-Aldrich, Ireland) supplemented with Protease Inhibitor cocktail 1 (1/100, Sigma-Aldrich, Ireland) and Phosphatase Inhibitors Cocktail 2 & 3 (1/500, Sigma-Aldrich, Ireland). Following lysis, total protein was quantified by BCA reagent and normalized (Thermo Fischer Scientific, Ireland). 20 μg of protein was resolved under reduced conditions by SDS-PAGE (Invitrogen, Ireland) and transferred to PVDF membrane (Thermo Fisher, Ireland) using a semi dry transfer system (Thermo Fisher Scientific, Ireland). Membranes were blocked for 1 h at room temperature in TBS-T (0.1%) - 5% milk (Sigma-Aldrich, Ireland). Primary antibodies were diluted in TBS-T (0.1%) - 5% BSA or milk (Table 2). Membranes were incubated with horseradish peroxidase conjugated anti-mouse or anti-rabbit antibodies (1/1000, R&D Systems, USA) in TBS-T (0.1%) 5% milk for 1 h at room temperature. Three 5 min washes in TBS-T (0.1%) were carried out pre- and post-secondary antibody incubation. Western blots were developed with enhanced chemiluminescence substrate (ThermoFisher Scientific, USA) using Amersham Imager (GE, USA).

**Table 2.** Antibodies used for Western Blot analysis

| **Antibody** | **Clone** | **Dilution** |
| --- | --- | --- |
| FKBPL | 10060-1-AP (Proteintech, USA) | 1/1000 |
| Phospho-p65<br>(Ser536) | 93H1 (Cell Signaling, USA) | 1/1000 |
| p65 | D14E12 XP (Cell Signaling, USA) | 1/1000 |
| IκBα | L35A5 (Cell Signaling, USA) | 1/1000 |
| Phospho-JNK<br>(Thr183/Tyr185) | G9 (Cell Signaling, USA) | 1/2000 |
| JNK | 9252 (Cell Signaling, USA) | 1/1000 |
| IL-1β | AF-401-NA (R&D Systems, USA) | 1/1000 |
| B actin | ab8227 (Abcam, USA) | 1/5000 |
| Aβ tubulin | 2148 (R&D Systems, USA) | 1/1000 |

### ELISA

For cytokine measurements, BMDMs were seeded at 5 × 10^5^ cell/ml in a 12-well plate. Cells were stimulated as indicated and supernatants removed and analyzed for Il-1β and TNF-α (both R&D Duoset ELISA kits) according to the manufacturer’s instructions. Cytokine measurements from *in vivo* peritoneal lavage washing (3 ml PBS) were analyzed for IL-6, TNF-α and IL-10 by ELISA (R&D Duoset ELISA kits).

### In vivo LPS survival

Mice were injected with 6 mg/kg *E. coli*-derived ultrapure LPS (Invivogen, UK) or PBS by intra perineal injection. Mice were treated with ALM201 (3 mg/kg), PBS or dexamethasone by sub cutaneous injection, as outlined in Supplementary Figure 1A. Mice were monitored over 60 h as per UK Home Office guidelines. Mice were culled immediately at a humane endpoint noted by loss of self-righting (Supplementary Figure 1B) and insensitivity to touch. Peritoneal lavage was conducted at endpoint to conduct cytokine analysis.

### Flow cytometry

LPS (6 mg/ml) and/or AD-01 (3 mg/kg) or equal volume sterile PBS was injected via intraperitoneal injection into C57BL/J6 wild-typeor *Fkbpl^+/−^* mice. Following a 3 h incubation mice were culled by cervical dislocation and peritoneal lavage was performed with 3 ml of PBS containing 3% BSA. Lavage fluid was harvested and cells were Fc blocked (Invitrogen, Ireland) and stained on ice for 30 min (Table 3) and gated as outlined in Supplementary Figure 2. Compensation was determined by single staining alone and no positive staining detected from IgG controls. Analysis was conducted using FlowJo Version 2 (Flowjo, USA).

**Table 3.**
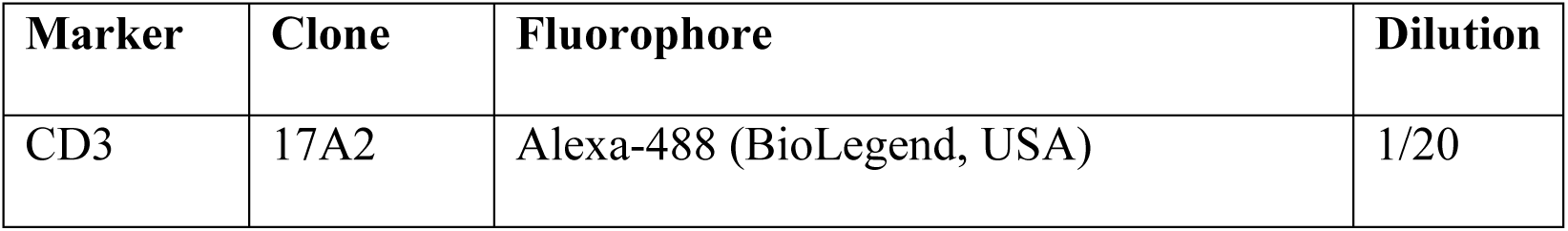
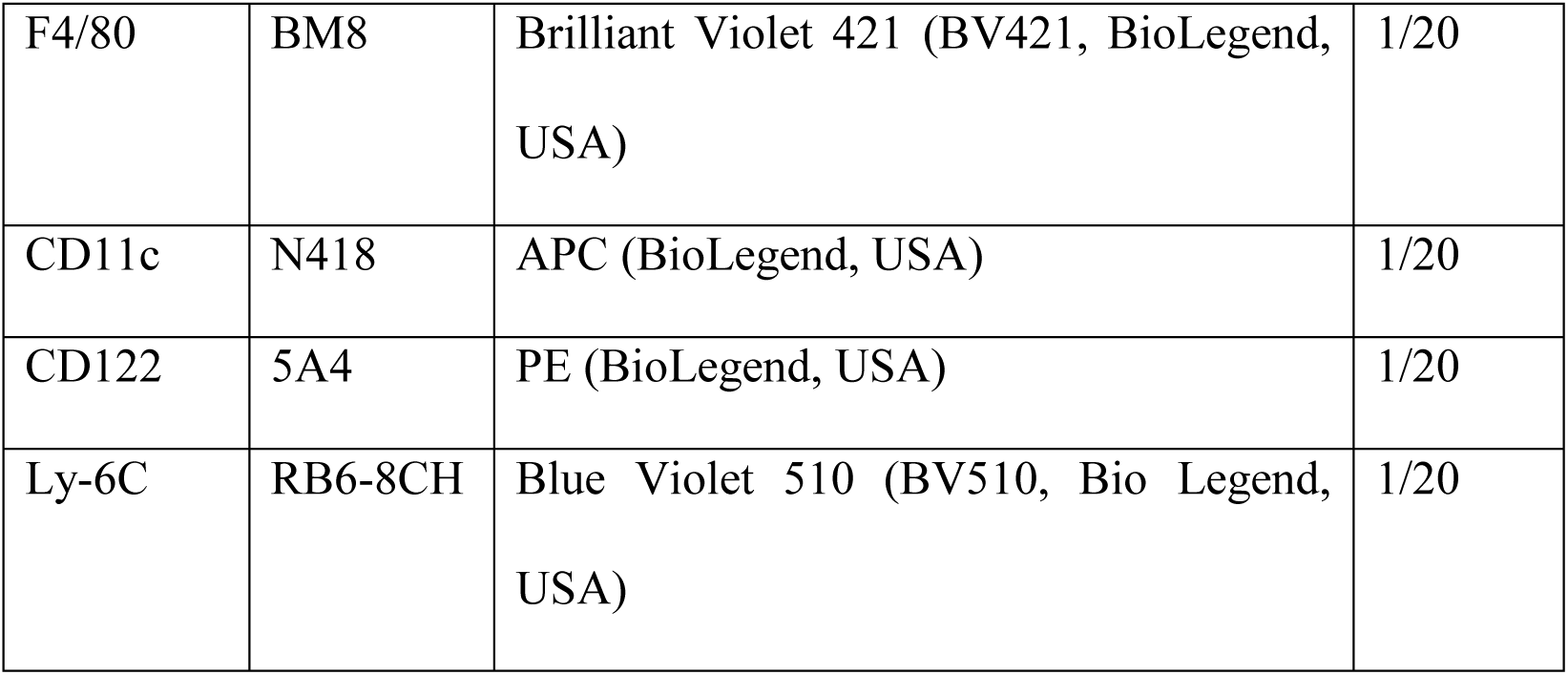
Antibodies used to characterise immune cell component

### In silico analysis

The global biobank engine (comprised of several biobanks from different geographical regions including Japan Biobank, UK Exome Biobank, Million Veterans Program; https://biobankengine.stanford.edu/) was used to assess the extent of variation in the human *fkbpl* gene, and whether variants found in *fkbpl, fkbp51* and *fkbp52* were associated with phenotypes of interest (tumor, autoimmune diseases, inflammatory diseases). A list of phenotypes which met the traditional genome-wide association study (GWAS) significance threshold of p<= 5×10^−8^ was compiled. The information obtained included the log fold change, average expression, P value and adjusted P value for each series (Supplementary Table 1).

The effect of genetic variation in *FKBPL* on tissue-specific gene expression levels was tested using the Genotype-Tissue Expression (GTEx) database (GTEx Consortium, 2014: PMID 23715323). Known variants in *fkbpl* were queried in the GTEx database, and any variants which were significantly associated (p<=5×10^−8^) with altered gene expression levels in a given tissue had their normalized expression levels (NES) in that tissue noted.

To investigate a link between *fkbpl* gene expression and psoriasis a search was performed on the NCBI GEO database to obtain mRNA microarray results and identified GSE14905 from skin biopsy samples (Affymetrix Human Genome U133 Plus 2.0 Array) which compared a non-disease control cohort and a diseased cohort. The dataset was downloaded from the NCBI GEO webpage via the R Studio program, and clinical data extracted to create .CEL files containing the study information. Using the Bioconductor Array Quality Metrics package on R Studio, quality control was undertaken for each data series. *Fkbpl* was defined as the gene of interest and using probe set 219187_at, each dataset was statistically analyzed to compare the expression of this gene in the control and diseased groups. Each boxplot showed the FKBPL gene expression profile of the two groups under investigation and any outliers in the analysis.

### Data Presentation and Statistical Analysis

All experimental data and statistical analysis were performed using the GraphPad Prism program (Graphpad Prism version 8.0 for Windows; GraphPad Software, Inc. San Diego, CA). Data is expressed as mean values ± standard error of the mean (SEM). To assess statistical differences, data was analyzed using Student’s 2-tailed t test or for comparison of multimer means an Anova was performed. For all statistical tests, P values <0.05 were considered significant.

## Results

### FKBPL is a novel regulator of LPS-induced endothelial permeability

Previous studies investigating the vasculature in *Fkbpl^+/−^* mice noted that they displayed enhanced angiogenesis with an associated increase in the number of erythrocytes leaking into surrounding tissues compared to wild-type mice (26). Therefore we hypothesized that FKBPL may be a novel regulator of vascular permeability. FKBPL knockdown, using siRNA transfection, resulted in ∼ 80% reduction in both mRNA and protein expression in human microvascular endothelial cells (HMEC-1); LPS stimulation did not affect the reduction in FKBPL mRNA or protein expression following transfection (Fig. 1A, Supplementary Fig. 3A). Next, we investigated if FKBPL regulated endothelial permeability using *in vitro* and *in vivo* Evan’s blue permeability assays. When FKBPL was knocked down in HMEC-1 cells there was an increase in barrier permeability compared to non-targeted control cells (Fig 1 B, p = 0.0347, n=3). We then investigated if *Fkbpl^+/−^* mice also displayed enhanced blood vessel permeability. Evan’s blue dye was injected intravenously and the dye was extracted from major organs. There was a significant increase in permeability in the skin of the FKBPL deficient mice (Fig. 1C. p=0.001378, n=3) which was visible externally (Fig. 1D). Endothelial barrier dysfunction occurs during stimulation by inflammatory agents or in inflammation-related disease states (40) and therefore we investigated if FKBPL had a role regulating LPS-induced endothelial permeability by utilizing the X-CELLigence system to measure cell impedence (41). Increasing doses of LPS resulted in a greater decrease in HMEC-1 cell impedance (Supplementary Fig. 3B) without affecting cell viability (Supplementary Fig. 3C). Real-time analysis demonstrated that FKBPL knockdown resulted in decreased impedance in HMEC-1s (Fig. 1E, F p=0.025, n=3) and this was further enhanced by LPS treatment (Fig 1E, F. p=0.0249, n=3). Analysis of gene expression showed that there was no change in the levels of the endothelial adhesion marker, VE cadherin, when FKBPL was knocked down (Fig. 1G). However, there was a significant upregulation of TNF following LPS stimulation and this was approximately three-fold higher when FKBPL was knocked down in the HMEC-1 cells (Fig. 1H, p=0.025, n=3). In addition, TNF mRNA was also increased three fold in unstimulated FKBPL-knockdown HMEC-1 cells compared to non-targeted controls (Fig. 1H; p<0.001, n=3). TNF upregulation is associated with activation of NF-κB activation and therefore we assessed phosphorylation of p65 in HMEC-1 cells following FKBPL knockdown. Indeed, we observed increased phosphorylation of p65 when FKBPL was knocked down in HMEC-1 cells following LPS stimulation (Fig. 1 I). Stimulation of FKBPL knockdown HMEC-1 cells with LPS resulted in a significant increase in the phosphorylation of p65 compared to non-targeted (NT) control cells at 90 min post treatment (Fig. 1 I; p=0.0367, n=3).

**Figure 1.**
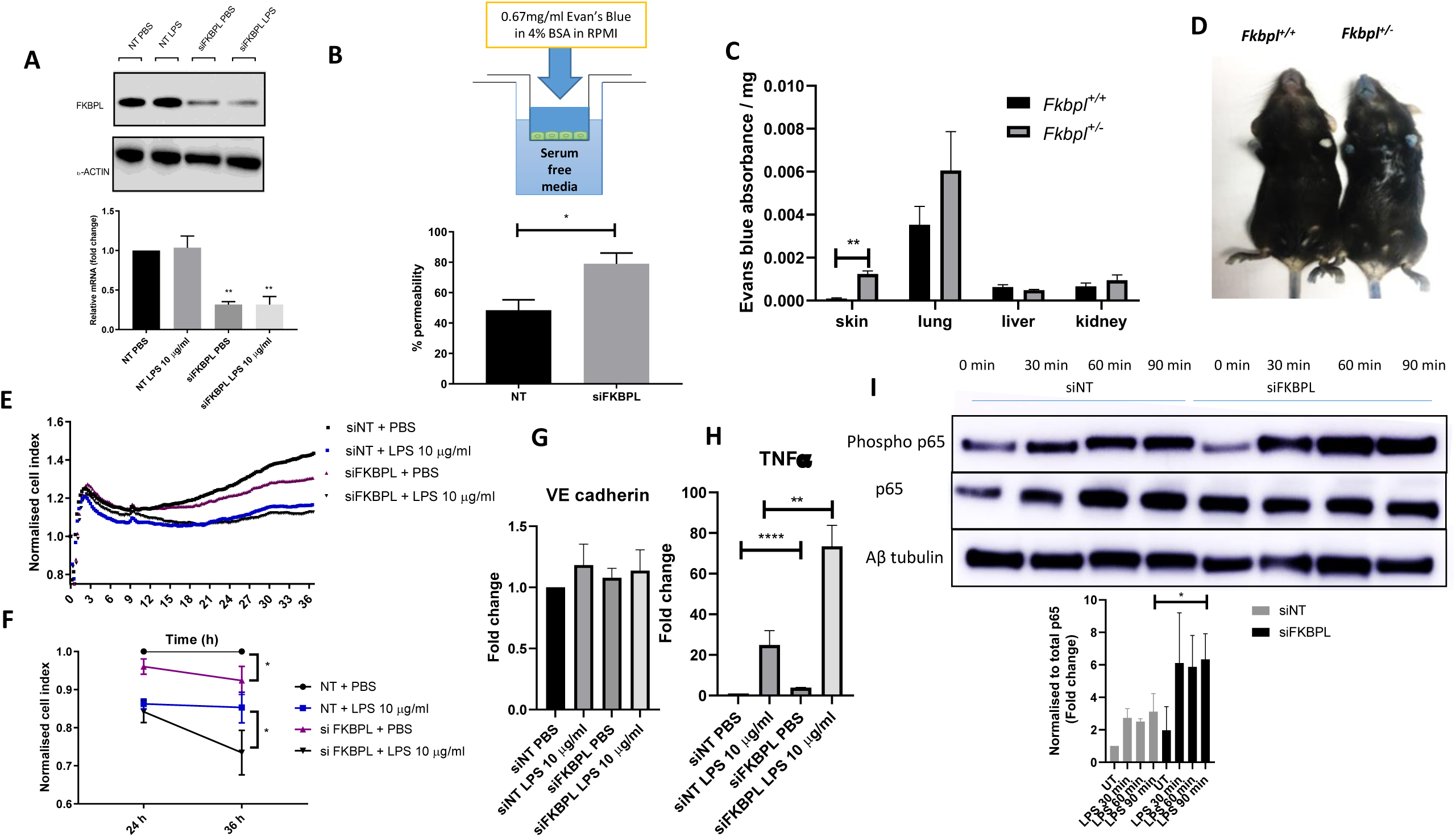
FKBPL is a novel regulator of endothelial permeability under normal and inflammatory conditions. **A)** FKBPL knockdown in human microvascular endothelial cells (HMEC-1s) using a siFKBPL and non-targeted (NT) control in untreated and LPS (10 μg/ml) stimulation conditions. FKBPL protein expression was analyzed by western blot and gene expression by qPCR. **(B)** *In vitro* Evan’s blue permeability assay in HMEC-1 cells transfected with siFKBPL or non-targeted (NT) control. **(C)** *In vivo* Evan’s blue permeability assay, Evan’s blue injected by tail view into *Fkbpl^+/−^* (haploinsufficient) or *Fkbpl^+/+^* (wild-type mice) and absorbance in major organs measured. **(D)** Representative images of the skin changes between *Fkbpl^+/−^* (haploinsufficient) or *Fkbpl^+/+^* (wild-type mice) following Evan’s blue intravenous injection. **(E)** Cell impedance was measured in HMEC-1s following transfection with siFKBPL or non-targeted (NT) control and treated with LPS (10 μg/ml) or PBS (control) for 36 h. **(F)** HMEC-1 cell index for each condition (siNT + PBS, siNT + LPS, siFKBPL+PBS, siFKBPL+LPS) was measured by the X-CELLigence system, normalised at either 24 or 36 h. Gene expression analysis of **(G)** VE cadherin and **(H)** TNF in HMEC-1 cells transfected with siFKBPL or non-targeted (NT) control and treated with PBS or LPS (10 μg/ml) by qPCR. **(I)** Protein expression of phosphorylation of p65 (Ser536), total p65 and αβ tubulin in HMEC-1 cells transfected with siFKBPL or non-targeted (NT) control and treated with PBS or LPS (10 μg/ml). Blot images represent one of three independent experiments. Protein expression was quantified using ImageJ, adjusted to αβ tubulin and normalized to control. Data points are mean ± SEM. n ≥ 3. * p < 0.05, ** p < 0.01, ***p < 0.01 (t-test)

### FKBPL based peptide, AD-01, abrogates LPS induced endothelial permeability

The FKBPL-based peptide, AD-01, has previously shown potent anti-angiogenic activity and therefore we assessed the ability of AD-01 to regulate endothelial permeability. Utilizing the X-CELLigence system, AD-01 treatment of HMEC-1 cells resulted in a small decrease in impedance, potentially indicating a change in cell morphology as previously described (26) (Fig. 2A, B p=0.0395). Next, we investigated if AD-01 could abrogate the decrease in cell impedance induced by LPS treatment. There was a significant increase in cell impedance upon addition of AD-01 indicating a potential protection of the endothelial barrier (Fig. 2C,D p=0.0346). To visualize the endothelial barrier, the HMEC-1 cells were sheared for 24 h before addition of LPS/AD-01 and then fixed and stained for VE-cadherin tight junctions. As expected, LPS induced a breakdown of VE cadherin junctions and this was abrogated in the presence of AD-01 (Fig. 2F, p<0.0001). There was no change in the gene expression of VE cadherin in HMEC-1 cells following LPS/AD-01 treatment suggesting that AD-01 caused a redistribution of VE cadherin at the cell surface (Fig. 1G).

**Figure 2.**
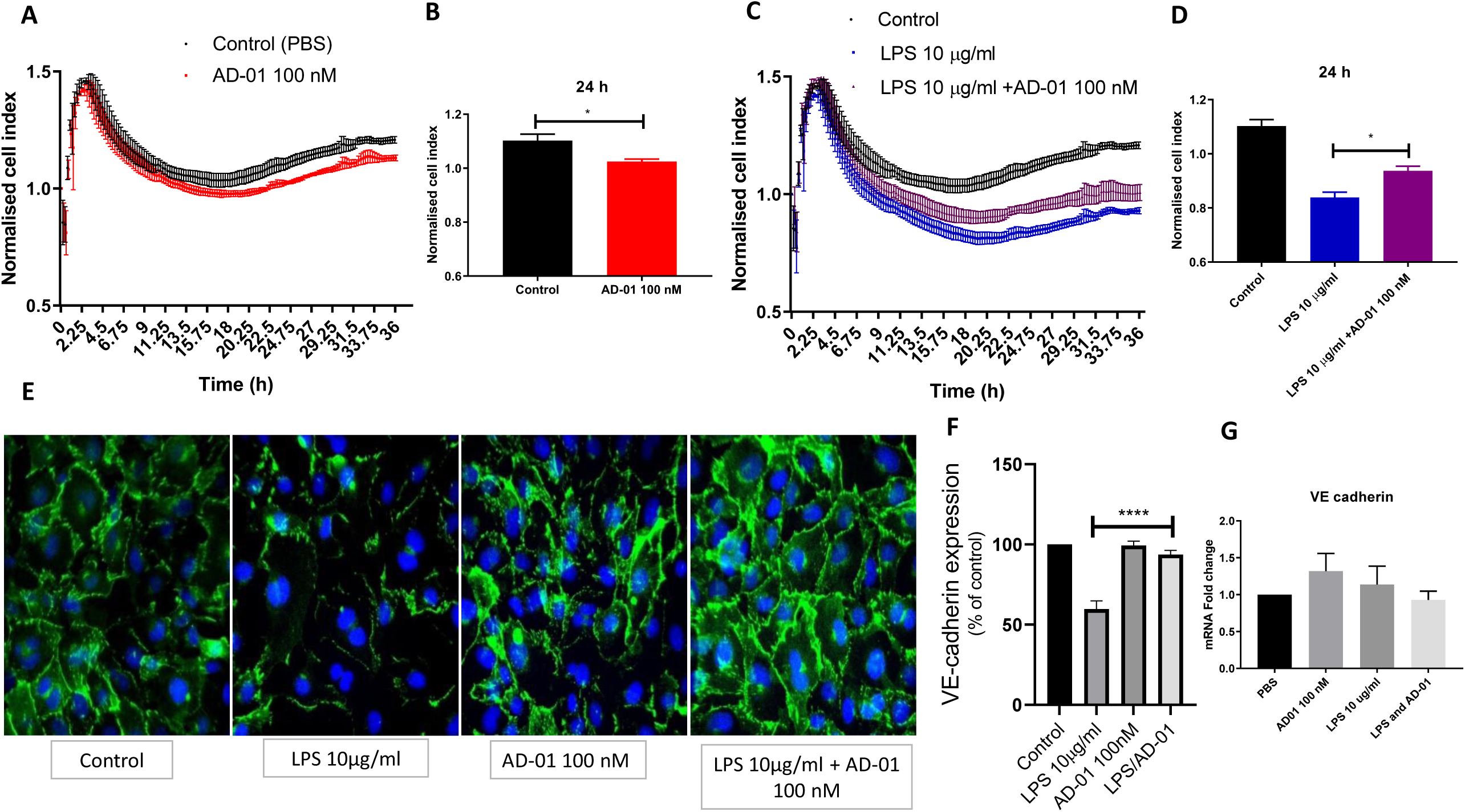
The FKBPL-based peptide, AD-01 promotes endothelial barrier function following LPS stimulation by stabilizing VE cadherin tight junction formation. **A)** Cell impedance was measured in human microvascular endothelial cells (HMEC-1s) following treatment with the FKBPL derived peptide AD-01(100 nM) or PBS (vehicle control) using the X-CELLigence system. **(B)** HMEC-1 cell index for each condition (PBS control or AD-01 100 nM) was measured by the X-CELLigence system and normalised to PBS control at the 24 h time point. **(C)** Cell impedance was measured in HMEC-1s following treatment with PBS (vehicle control), LPS (10 μg/ml) alone or LPS (10 μg/ml) and FKBPL derived peptide AD-01(100 nM) using the X-CELLigence system. **(D)** HMEC-1 cell index for each condition (PBS, LPS + AD-01 100 nM), measured by the X-CELLigence system, and normalized to PBS control at the 24 h time point. **(E)** HMEC-1s were cultured under shear stress conditions to enhance tight junction formation. Following treatment (PBS, AD-01, LPS, LPS+AD-01) for 2 h, VE cadherin tight junctions were visualized using immunofluorescence and representative images displayed of 4 independent experiments displayed. **(F)** Quantification of shear stressed HMEC-1s VE cadherin expression treated with PBS, AD-01, LPS and LPS+AD-01 for 2 h **(G)** *VE cadherin* gene expression of HMEC-1s treated with PBS (control), AD-01 (100 nM), LPS (10 μg/ml), LPS+AD-01) by qPCR. Data points are mean ± SEM. n ≥ 3. * p < 0.05, ** p < 0.01, ***p < 0.01 (t-test or one way ANOVA).

### FKBPL and its pre-clinical peptide, AD-01, decrease phosphorylation of p65 following LPS stimulation in macrophages by a non CD44-dependent mechanism

NF-κB signaling is crucial for the inflammatory response by innate immune cells and therefore we investigated if FKBPL could also regulate activation of NF-κB activation in macrophages (42). BMDMs from wild-type and *Fkbpl^+/−^* mice were isolated and stimulated with LPS (100 ng/ml). Similar to microvascular endothelial cells, *Fkbpl^+^*^/-^ BMDMs exhibited increased phosphorylation of p65 following LPS stimulation (Fig. 3A, 60 min p=0.037, n=3). Next wild-type BMDMs were stimulated with LPS (100 ng/ml) ± AD-01 (1 nM). The addition of AD-01 abrogated phosphorylation of p65 following LPS stimulation in the presence of ATP (Fig. 3B 45 min p = 0.0426; 60 min p=0.003; n=3). FKBPL and AD-01 have been shown to exert anti-angiogenic activity through the cell surface receptor CD44 (26). To investigate if the ability of AD-01 to regulate NF-κB signaling is dependent upon CD44, we used a CD44 knockout mouse model. BMDMs were extracted from CD44 knockout mice (CD44^−/−^) and stimulated with LPS (100 ng/ml) ± AD-01 (1 nM). AD-01 also abrogated phosphorylation of p65 in CD44^−/−^ BMDMs, thus indicating that FKBPL regulates NF-κB activation independently of CD44 (Fig. 3C 90 min p= 0.0295; n=3). LPS can also activate other intracellular signaling events downstream of Toll-like receptor 4 (TLR4) including the c-Jun N-terminal kinase (JNK) pathway (43). However, AD-01 did not modulate phosphorylation of JNK (Fig. 3D).

**Figure 3.**
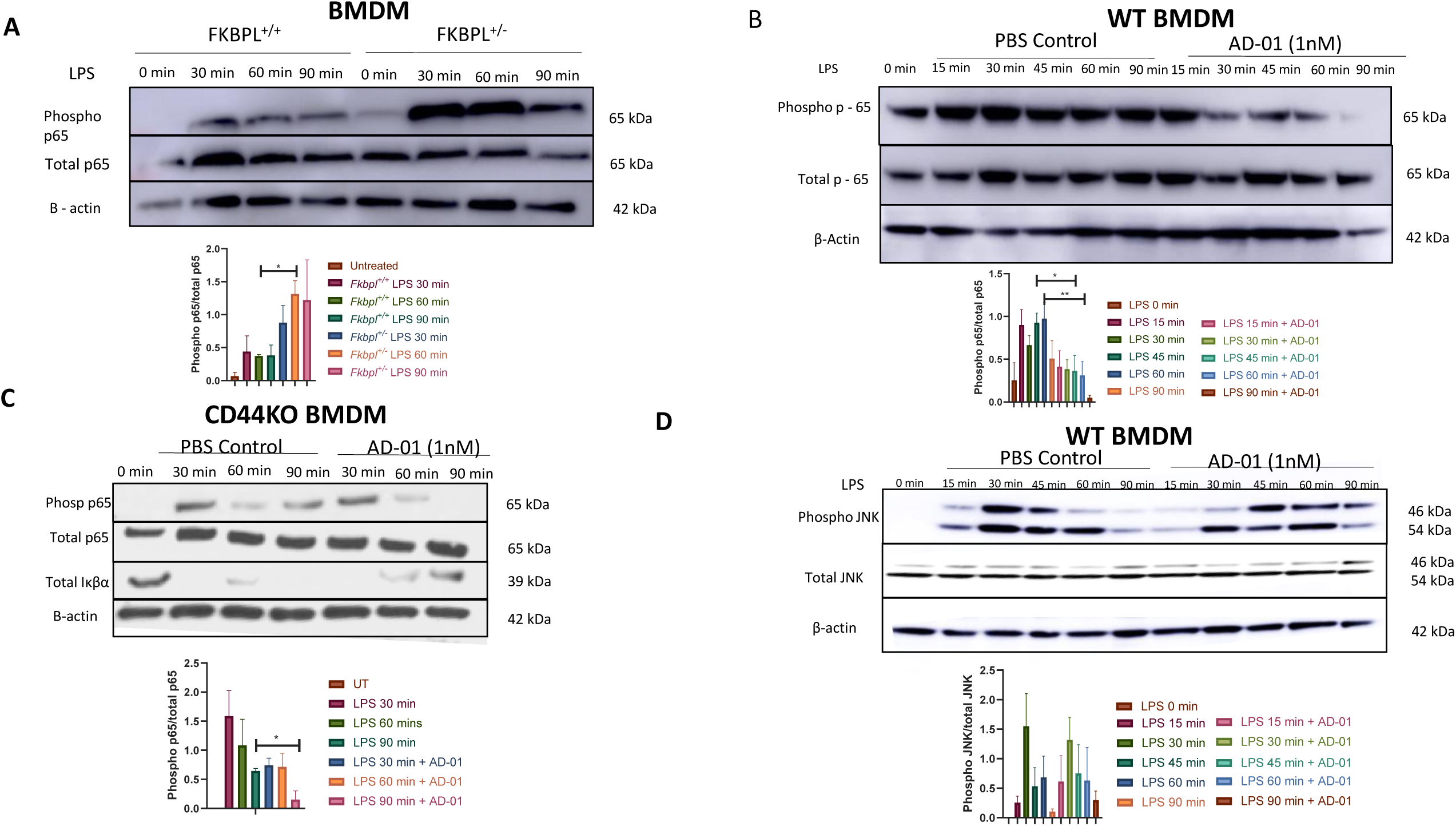
FKBPL and its peptide derivatives are negative regulators of NF-κB activation. **(A)** Bone marrow derived macrophages (BMDMs) from *Fkbpl^+/−^* (haploinsufficient) and *Fkbpl^+/+^* (wild-type mice) where isolated and treated with LPS for 30, 60, 90 min. Protein was extracted and NF-κB activation measured by Western blot analysis of phosphorylation of p65 (Ser536). **(B)** BMDMs from wild-type mice were treated with LPS (100 ng/ml) or LPS + AD-01 (1 nM) for 15, 30, 45, 60, 90 min and protein extracted. Phosphorylation of p65(Ser536) was analyzed by Western blot. **(C)** BMDMs from CD44 knockout and wild-type mice where isolated and treated with LPS for 30, 60, 90 min. Protein was extracted and NF-κB activation measured by Western blot analysis of phosphorylation of p65 (Ser536) and Iκβα expression. **(D)** BMDMs from wild-type mice were treated with LPS (100 ng/ml) or LPS + AD-01 (1 nM) for 15, 30, 45, 60, 90 min and protein extracted. Phosphorylation of JNK (Thr183/Thr185) was analyzed by Western blot. Blot images represent one of three independent experiments.

### FKBPL based peptide, AD-01, decreases gene expression and secretion of proinflammatory mediators in macrophages following LPS stimulation

NF-κB activation induces the expression and secretion of proinflammatory cytokines. The ability of AD-01 to inhibit the secretion of TNF and IL-1β in BMDMs was investigated using ELISA. BMDMs from wild-type and CD44^−/−^ mice were stimulated for 6 h with LPS (100 ng/ml) ±AD-01 (1 nM and 100 nM) and supernatants collected. There was a significant ∼ 25% decrease in TNF secretion following AD-01 (1 nM) treatment in both the wild-type BMDMs (Fig. 4A, p=0.0387; n=3) and CD44^−/−^ BMDMs (Fig.4 A, p=0.0380; n=3). The production of bioactive IL-1β requires two sequential steps involving priming and activation of the NLRP3 inflammasome. Priming entails the recognition of danger-associated molecular patterns or pathogen-associated molecular patterns, such as LPS, by TLR4, thereby resulting in activation of NF-κB and induction of pro-IL1β. Subsequently NLRP3 agonists (such as ATP) results in the formation of the NLRP3 inflammasome complex and pro-IL-1β is cleaved to its mature form and secreted (44). BMDMs were treated with LPS (100 ng/ml) for 3 h, then PBS control or AD-01 (1 nM) were added for 1 h in serum free media and then ATP (5 nM) for 45 min. There was a significant reduction in IL-1β secretion in the AD-01 treated BMDMs following addition of ATP (Fig. 4B p=0.0086; n=4). Furthermore, there was no change in expression of pro-IL1-β expression in the AD-01 treated group following ATP, indicating that the reduction in IL-1β secretion was due to an inhibitory effect on the NF-κB priming of the inflammasome (Fig. 4C). There was also a significant reduction in TNF secretion in the AD-01 treated cells following addition of LPS and LPS and ATP (Fig. 4D; LPS+AD-01 p=0.0054, LPS+ATP+AD-01 p =0.0134; n=5). The expression of NF-κB regulated genes following treatment with LPS (100 ng/ml) and rFKBPL (50 ng/ml) or AD-01 (1nM) was investigated using qPCR. There was a significant reduction in IL-1β expression (AD-01, p=0.0314; n=6), IL-18 expression (rFKBPL p=0.0292; n=6); NFKB1 (AD-01, p=0.0239 rFKBPL, p=0.0185; n=5), COX2 (AD-01, p=0.0064 rFKBPL, p=0.0185; n=5) and IL-6 (rFKBPL, p=0.0439; n=5) (Fig. 4E.). Similarly, in THP-1 human monocyte cell line, AD-01 (1 nM) significantly reduced mRNA expression of *tnf*, *Il-β*, *Il-18*, *nlrp3*, *nfkb1* and *casp1* following 6 h stimulation with LPS (1000 ng/ml) (Supplementary Fig. 4).

**Figure 4.**
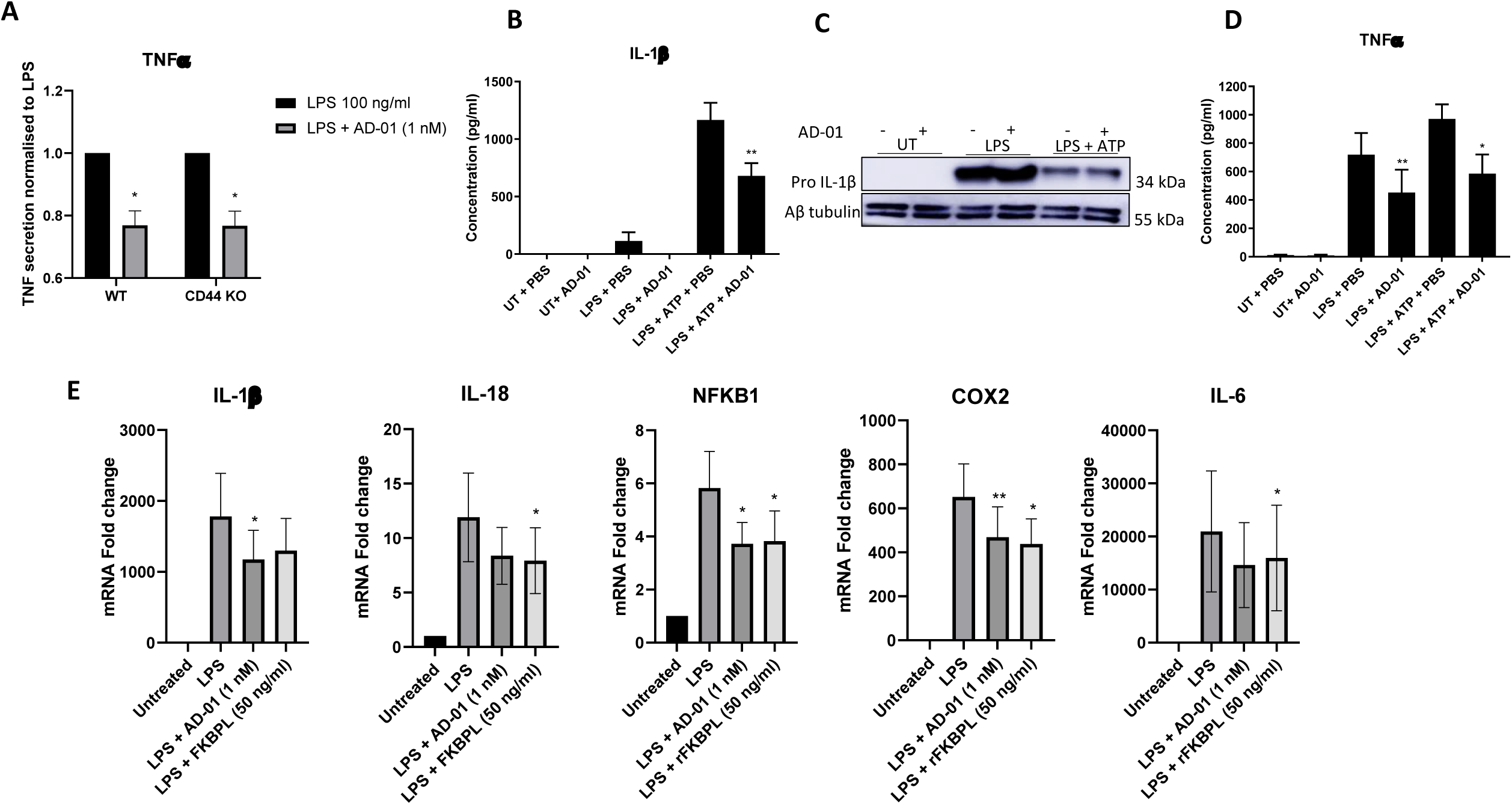
FKBPL and its peptide derivative, AD-01, decreases proinflammatory gene expression and secretion following LPS stimulation. **(A)** BMDMs from wild-type and CD44-knockout mice were stimulated with LPS (100 ng/ml) and treated with PBS (vehicle control) or AD-01 (1 nM). Supernatant was collected after 6 h and secretion of TNF quantified by ELISA. **(B)** BMDMs were treated with LPS (100 ng/ml) for 3 h then media replaced with serum free media. PBS or AD-01 (1 nM) was added for 1 h and then ATP (5 nM) to stimulate IL-1β secretion. IL-1β was detected by ELISA for each treatment condition. **(C)** Pro-IL-1β expression was detected by western blot and **(D)** TNF secretion was detected by ELISA. **(E)** BMDMs were stimulated with LPS (100 ng/ml) and treated with PBS (vehicle control), AD-01 (1 nM) or rFKBPL (50 ng/ml) for 6 h. NF-κB regulated genes (*IL-1β, IL-18, NFKB1, COX2, IL-6*) were analyzed by qPCR. Data points are mean ± SEM. n ≥ 3. * p < 0.05, ** p < 0.01, ***p < 0.01 (t-test or one way ANOVA). Blot image represent 1 of 3 independent experiments.

### FKBPL and its clinical peptide, ALM201, are protective against LPS lethality *in vivo*

To investigate the protective role of FKBPL *in vivo*, the LPS survival model was utilized. *Fkbpl^+/−^* mice had a significantly shorter survival than wild-type mice (Fig. 5A, p=0.0021, n>5 mice/group). The clinical peptide based on FKBPL, ALM201, has enhanced stability to AD-01 and was utilized in this *in vivo* model. Briefly, group 1 mice received PBS as a control, group 2 mice received LPS (6 mg/kg), group 3 mice received ALM201 (3 mg/kg) prior to LPS and 3 more doses of ALM201 thereafter, group 4 mice received three doses of ALM201 (3 mg/kg) after LPS and group 5 mice received three doses of dexamethasone (10 μg) (Supplementary Fig. 1A). As expected, the LPS only group had a significant shorter survival than the PBS treated group (Fig. 5B, C; p = 0.0018, n=5/group). Furthermore, dexamethasone significantly increased survival compared to LPS alone (Fig. 5B, C; p=0.0044, n=5/group). Pre-treatment with ALM201 resulted in a comparable significant increase in survival to the dexamethasone treated group (Fig. 5B, C; p=0.0342; n=5/group). Impressively, treatment of ALM201 post LPS resulted in 100% survival (Fig. 5D; p=0.0009, n=5/group). Analysis of cytokine concentrations following peritoneal lavage at experimental endpoint revealed that ALM201 post treatment significantly reduced IL-6 (Fig. 5E; p<0.0001) and TNF (Fig. 5F: p<0.0001) pro-inflammatory cytokines compared to LPS only and significantly increased the anti-inflammatory cytokine, IL-10 (Fig. 5G; p=0.004). Finally, we investigated if FKBPL and its peptide derivatives had an effect on *in vivo* inflammatory cell infiltrates. Wild-type and *Fkbpl^+/−^* mice were injected with intra peritoneal LPS (6 mg/kg) and PBS (vehicle control) or AD-01 (3 mg/kg) for 3 h and peritoneal lavage washings were collected. LPS treatment resulted in a significant decrease in peritoneal macrophages and γδ T cells and a non-significant increase in neutrophils (Fig. 5H, Supplementary Figure 2B). However, there was no significant changes in macrophages, neutrophils or γδ T cells between wild-type and *Fkbpl^+/−^* mice or following AD-01 treatment (Fig. 5H, Supplementary Figure 2B). Furthermore, there was no change in NK cells or T cells following LPS injection (Supplementary Fig. 2B).

**Figure 5.**
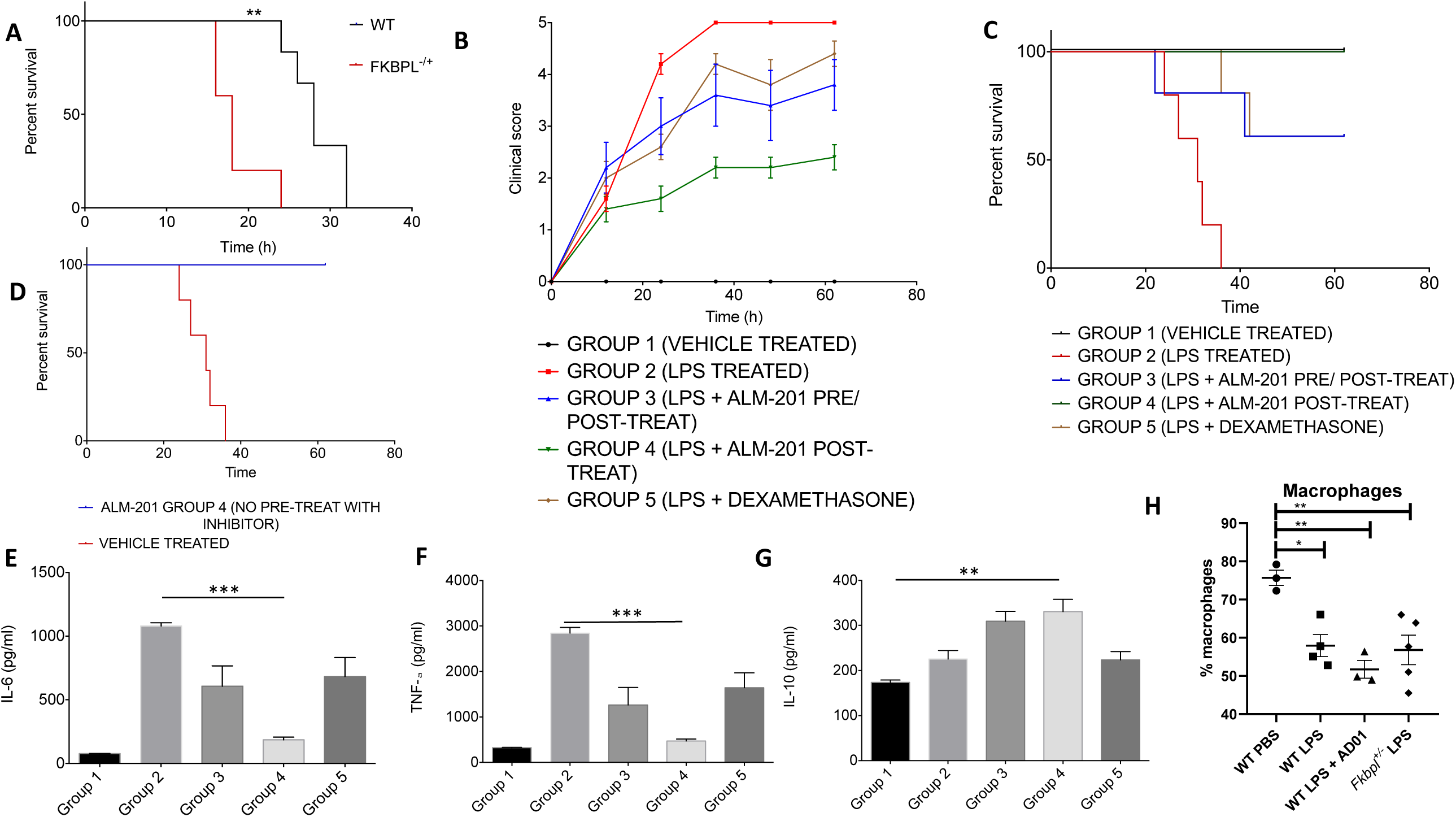
The FKBPL based clinical peptide, ALM201, abrogates *in vivo* LPS induced lethality. **(A)** *Fkbpl^+/−^* (haploinsufficient) and *Fkbpl^+/+^* (wild-type mice) where treated with intraperitoneal LPS (6 mg/kg) and monitored for 60 h. Mice were sacrificed upon reaching the humane endpoint. **(B)** C57BL/6 mice following treatment with PBS control (Group 1), LPS 6 mg/kg (Group 2), LPS + AD-01 3 mg/kg pre and post LPS (Group 3), LPS + AD-01 3 mg/kg post LPS (Group 4), LPS + dexamethasone 10 μg per mouse post LPS (Group 5). Clinical scoring using the following criteria: score 0, no symptoms; score 1, piloerection and huddling; score 2, piloerection, huddling and diarrhea; score 3, lack of interest in surrounds and severe diarrhea; score 4 decreased movement and listless appearance; score 5, loss of self-righting reflex. **(C)** Survival analysis was monitored for 50 h and mice were humanely scarified upon reaching clinical score 5**. (D)** Survival analysis of LPS treated mice (Group 2) and LPS and ALM201 post treated mice (Group 4). Statistical significance of survival analysis was assessed by Kaplan-Meier log rank χ^2^ test. At experimental endpoint, peritoneal lavage washings were collected and assayed for **(E)** IL-6, **(F)** TNF **(G)** IL-10 by ELISA. **(H)** Wild-type mice treated with PBS, LPS 6 mg/kg and AD-01 3 mg/kg (post LPS) and *Fkbpl^+/−^* mice treated with LPS 6mg/ml were humanly sacrificed after 3 h and peritoneal lavage washings collected. The percentage of macrophages present was assessed by flow cytometry. Data points are mean ± SEM. n ≥ 3. * p < 0.05, ** p < 0.01, ***p < 0.01 (one way ANOVA).

### *In silico* analysis of FKBPL and inflammatory disorders

In order to determine if genetic variations in FKBPL are linked to human traits or diseases, we utilized data from the UK Biobank. The strongest genotype-phenotype association was a C/T missense SNP (rs28732176) which was strongly associated with psoriasis (p=1.78 x 10^−210^) (Fig. 6A). In addition, the rs28732176 variant was also significantly associated with rheumatoid arthritis (p= 5.58 x 10^−8^) (Fig. 6A). Furthermore, an A/G variant in the 5’ untranslated region of FKBPL (rs204892) was significantly associated with rheumatoid arthritis (p=7.84 x 10^−11^) as well as hay fever rhinitis or eczema (p=5.29 x 10 ^−13^) (Fig. 6A). Finally, a G/C missense variant SNP (rs35580488) was associated with lymphocyte count (p=1.48×10^−11^) (Fig. 6A). Interestingly, this variant is within the region of the FKBPL-based therapeutic peptides, AD-01 and ALM201, and this allele frequency is associated with approximately 1% of the population (Fig. 6B). As FKBPL shares similar homology to FKBP51 and FKBP52, particularly in the TPR domains, we also investigated if similar traits were associated with FKBP51 and FKBP52. FKBP51 had several variants associated with lymphocyte count but neither FKBP51 nor FKBP52 were associated with psoriasis, rheumatoid arthritis or any other disease with an inflammatory component (Supplementary Table 1). We used molecular modelling to predict any potential changes in structure from the missense SNPs, rs28732176 and rs35580488 using the Robetta server (45). The corresponding mutations, A90T and T46R/M could potentially influence the conformational flexibility of the PPIase domain and impact interactions with various cellular binding partners (Fig. 6C). Next, we investigated if reported SNPs in *fkbpl* had any predicted effect on gene expression. Indeed, rs28732176 (associated with psoriasis) is predicted to reduce *fkbpl* gene expression in both sun exposed, lower leg skin (Supplementary Table 2; NES = −0.24; p = 1.3×10^−13^) and non-sun exposed suprapubic skin (Supplementary Table 2; NES = −0.32; p= 1.8×10^−12^). Predicted gene changes associated with rs35580488 and rs204892 are presented in Supplementary Table 3 and 4. We then utilized the publicly available gene expression data series GSE14905 to further investigate *fkbpl* expression in psoriasis. *fkbpl* gene expression was downregulated in the lesional samples of psoriasis patients compared to healthy skin biopsies (Fig. 6C; p=3.19×10^−13^). Furthermore, *fkbpl* expression was also decreased in the lesional skin compared to the non lesional skin within diseased patients (Fig 6D; p=7.86×10^−15^).

**Figure 6.**
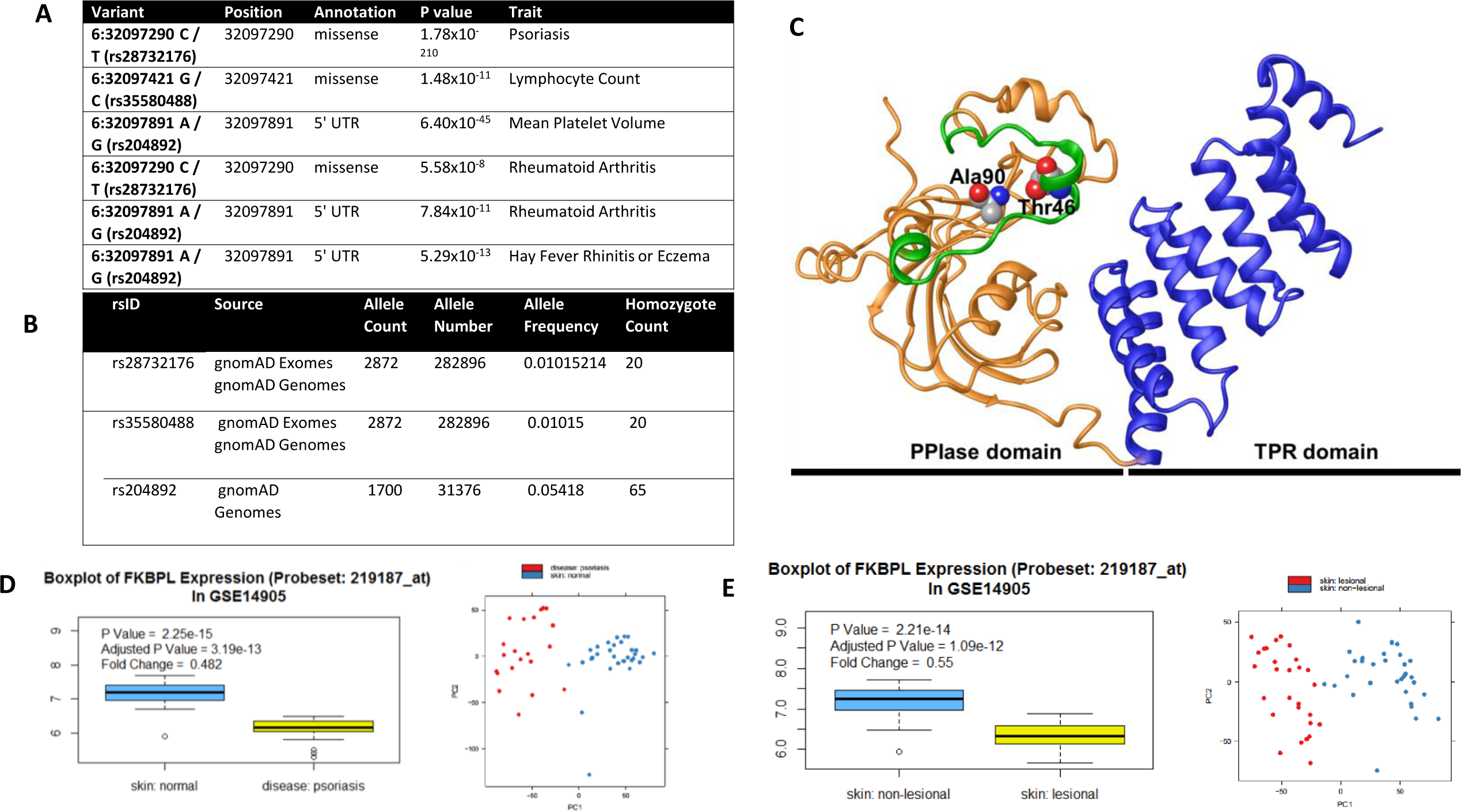
Single nucleotide polymorphisms in FKBPL are associated with auto-inflammatory disorders. **(A)** Single nucleotide polymorphism (SNPs) in FKBPL is associated with autoimmune disorders in the UK biobank dataset. **(B)** Frequency of identified SNPs in the general population within the UK biobank cohort. **(C)** A computational 3D-model of FKBPL, built based on FKBP51 using the Robetta server. The PPIase and TPR domains of FKBPL are shown in orange and blue. The ALM201 peptide is highlighted in green. Residues Ala90 and Thr46 that have mutations associated with inflammatory disorders are shown in a space-filling model. **(D)** Box plot comparing *fkbpl* gene expression in the normal skin biopsies of control samples and the psoriasis lesional skin biopsies of diseases samples in the GSE14905 dataset. **(E)** Box plot comparing FKBPL gene expression in the non-lesional and lesional skin biopsies of psoriasis patients in the GSE14905 dataset.

## Discussion

Here, for the first time, we describe a novel role for FKBPL as a negative regulator of NF-κB activation. Peptides based on the N terminal region of FKBPL prevent activation of NF-κB following LPS stimulation resulting in promotion of tight junctions in endothelial cells (Fig. 2) and decreased phosphorylation of p65 and cytokine release from macrophages (Fig. 3,4). Impressively, the clinical FKBPL peptide, ALM201, also abrogated LPS induced death *in vivo* (Fig. 5); whilst *Fkbpl^+/−^* mice demonstrated reduced survival to LPS. NF-κB has long been an attractive drug target for inflammation, however, the toxicity associated with indiscriminate blockade of NF-κB signaling has hindered clinical development (12, 46). ALM201 has completed a Phase 1a dose escalation clinical trial in the cancer setting and encouragingly, ALM201 had a favorable pharmacokinetic and safety profile, with no major adverse events noted (33). This study describes novel indications for FKBPL based therapeutics as anti-inflammatory agents via modulation of NF-κB signaling in endothelial cells and macrophages.

We had previously observed that *Fkbpl^+/−^* mice appeared to have less robust blood vessels compared to wild-type mice, despite enhanced angiogenesis (27). Indeed, here we demonstrate, using *in vitro* and *in vivo* Evan’s blue-based vascular permeability assays, that knockdown of FKBPL in endothelial cells decreases vascular integrity (Fig. 1). A reduction in VE-cadherin at endothelial cell junctions induced by pro-inflammatory factors, such as cytokines and LPS, results in increased permeability allowing inflammatory cells to migrate from the blood to tissues, as well as, exposure of the basement membrane and activation of the clotting cascade (47, 48). We show that addition of LPS to endothelial cells deficient in FKBPL results in enhanced LPS induced endothelial permeability (Fig. 1E, F) coupled with enhanced *tnf* gene expression (Fig. 1H) and increased phosphorylation of p65 (Fig. 1I). However, addition of AD-01 to LPS stimulated microvascular endothelial cells partially abrogated the decrease in impedence (Fig. 2) and enhanced VE cadherin at endothelial cell junctions (Fig. 2); which would be advantageous in some proinflammatory disease settings. FKBP51, another member of the FKBP protein family with TPR domains, has also been associated with regulation of endothelial barrier permeability, although it has not been established if FKBP51 can protect the endothelial under inflammatory conditions (49, 50). The underlying basic mechanisms of vascular integrity remain incompletely understood and to date this is the first report of an FKBP protein regulating endothelial permeability during the inflammatory response.

Next we investigated if FKBPL also had a role in modulating NF-κB signaling in macrophages. BMDMs derived from *Fkbpl^+/−^* mice exhibited enhanced phosphorylation of p65 compared to wild-type BMDMs when stimulated with LPS (Fig. 3A). Furthermore, treatment with AD-01 inhibited phosphorylation of p65 in wild-type BMDMs stimulated with LPS (Fig. 3B). AD-01 has previously been shown to require the cell surface receptor CD44 to inhibit angiogenesis and we hypothesized that CD44 may be involved in AD-01’s anti-inflammatory activity (26). LPS signals through TLR4, and CD44 has been shown to associate with TLR4 in macrophages and down regulate its signaling (51–53). Treatment with AD-01 also inhibited the phosphorylation of p65 and reduced secretion of TNF in the CD44^−/−^ BMDMs, indicating that CD44 is not required for FKBPL to modulate NF-κB signaling (Fig. 3C, 4A). Other members of the FKBP protein family with a similar structure to FKBPL have been shown to regulate NF-κB signaling (17). There is good sequence conservation between FKBPL and FKBP51 and FKBP52 in terms of both quantity and configuration of the TPR domains, whilst there is weak conservation in the N terminal regions which contains functional or non-functional PPIase domains of FKBPL, FKBP51 and FKBP52 (20). FKBP51 complexes with cytoplasmic p65 in unstimulated cells, and upon stimulation, FKBP51 is exchanged for FKBP52 and FKBP52 is then recruited to the promoter region of NF-κB genes (14, 17). The TPR domains of FKBP51 and FKBP52 are not required for the regulation of NF-κB signaling whilst the PPIase activity of FKBP52 is required for its NF-κB stimulatory activity, whereas the PPIase activity is not required for FKBP51 inhibitory action (14, 17). Similarly, FKBPL peptides AD-01 and ALM201 are based on the N terminal region of FKBPL which contains the non-functional PPIase domain (20). As AD-01 has similar activity to recombinant full length FKBPL in regulating NF-κB target genes, it is likely this is also the domain in FKBPL that is important for modulation of NK-κB (Fig. 4E). Further work is required to understand the intricate relationships between FKBPL, FKBPL51 and FKBP52 in the modulation of NF-κB signaling.

The data presented in this manuscript indicates that FKBPL-based therapeutics have potential utility as dual vascular stabilization and anti-inflammatory agents in a wide range of disorders. Decreased vascular integrity is a hallmark of serious pathological conditions and there are currently no therapies for stabilizing the vasculature (54–58). In addition, the systemic inflammatory response and production of pro-inflammatory cytokines further perpetuates the multiple organ dysfunction experienced in these patients (59). Although the use of glucocorticoids in sepsis is not conclusive, the RECOVERY trial showed dexamethasone treatment resulted in lower mortality in COVID patients receiving respiratory support (60, 61). Notably, in the *in vivo* LPS survival model, ALM201 therapy demonstrated enhanced survival compared to dexamethasone treatment (Fig. 5C), potentially indicating more potent activity. Considering that FKBPL based peptides have already been shown to be non-toxic in clinical and preclinical studies, this data indicates they may be repurposed as much needed therapeutics for these clinical indications with minimal risk of adverse effects.

Finally, we conducted *in silco* analysis for phenotypic traits associated with SNPs in FKBPL. Several SNPs in FKBPL were identified and associated with chronic diseases with autoimmune and endothelial dysfunction pathogenic traits. Psoriasis is a skin disease with a worldwide prevalence of 2% and it has a strong genetic predisposition (62). A missense SNP in FKBPL displayed a very strong association with psoriasis and furthermore, FKBPL gene expression was significantly downregulated in psoriatic skin lesions compared to non-lesional skin (Fig. 6). Anti-inflammatory biological therapies have shown good clinical efficacy in psoriasis patients. However, in some cases an initial clinical response is short lived and therapy resistance develops (62, 63). In addition to enhanced inflammation, the pathophysiology of psoriasis shows a strong pro-angiogenesis and endothelial dysfunction component (6, 64). Therefore, FKBPL based therapies could potentially have a therapeutic advantage by dual-targeting the dysfunctional vasculature as well as the inflammatory component of psoriasis lesions. Similarly, SNPs in FKBPL were associated with rheumatoid arthritis (Fig. 6), another chronic autoimmune disorder associated with both aberrant angiogenesis, endothelial dysfunction and inflammation and FKBPL may be a novel therapeutic that could be harnessed in this setting (65, 66). Of particular note is rs28732176, a missense mutation, associated with both psoriasis and rheumatoid arthritis (Fig. 6A). Interestingly it is within the region of FKBPL where the anti-inflammatory peptides, AD-01 and ALM201 are based and this variant is prevalent in an estimated 1% of the population (Fig. 6B).

In summary, for the first time we have shown that FKBPL is a novel regulator of endothelial permeability and pro-inflammatory cytokine signaling in response to inflammation. Furthermore, FKBPL-based peptides are potent inhibitors of endothelial barrier dysfunction and inflammatory cytokine signaling through modulation of NF-κB signaling using *in vitro* and *in vivo* models. We have also demonstrated that SNPs in FKBPL are associated with inflammatory conditions and suggest that FKBPL-based therapies may offer a novel therapeutic strategy for treatment of both acute and chronic inflammatory disorders.

---

**Supplementary Figure 1.**
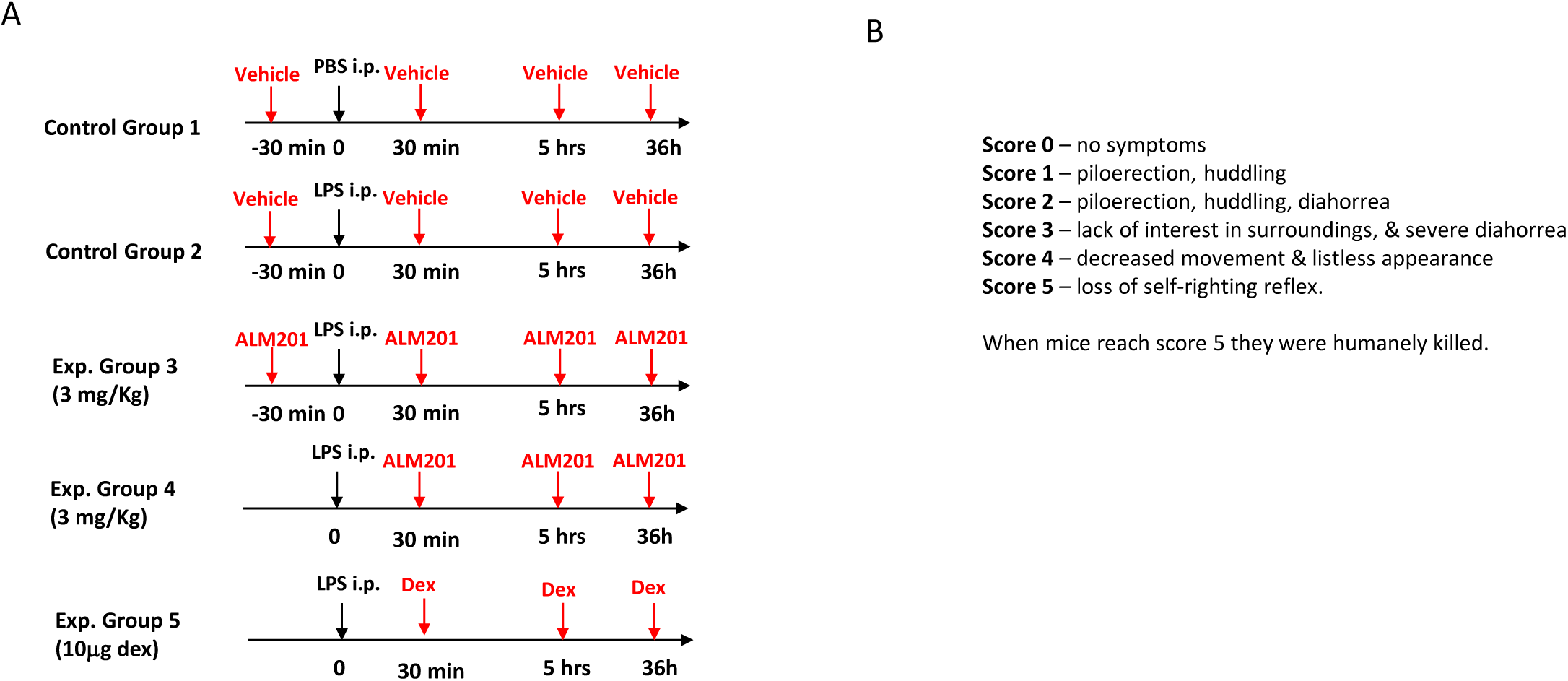
**A)** Schematic diagram displaying the treatment schedule of C57BL/6 mice with PBS control (Group 1), LPS 6 mg/kg (Group 2), LPS + AD-01 3 mg/kg pre and post LPS (Group 3), LPS + AD-01 3 mg/kg post LPS (Group 4), LPS + dexamethasone 10 μg per mouse post LPS (Group 5). **(B)** Clinical scoring of mice used for *in vivo* LPS survival experiments.

**Supplementary Figure 2.**
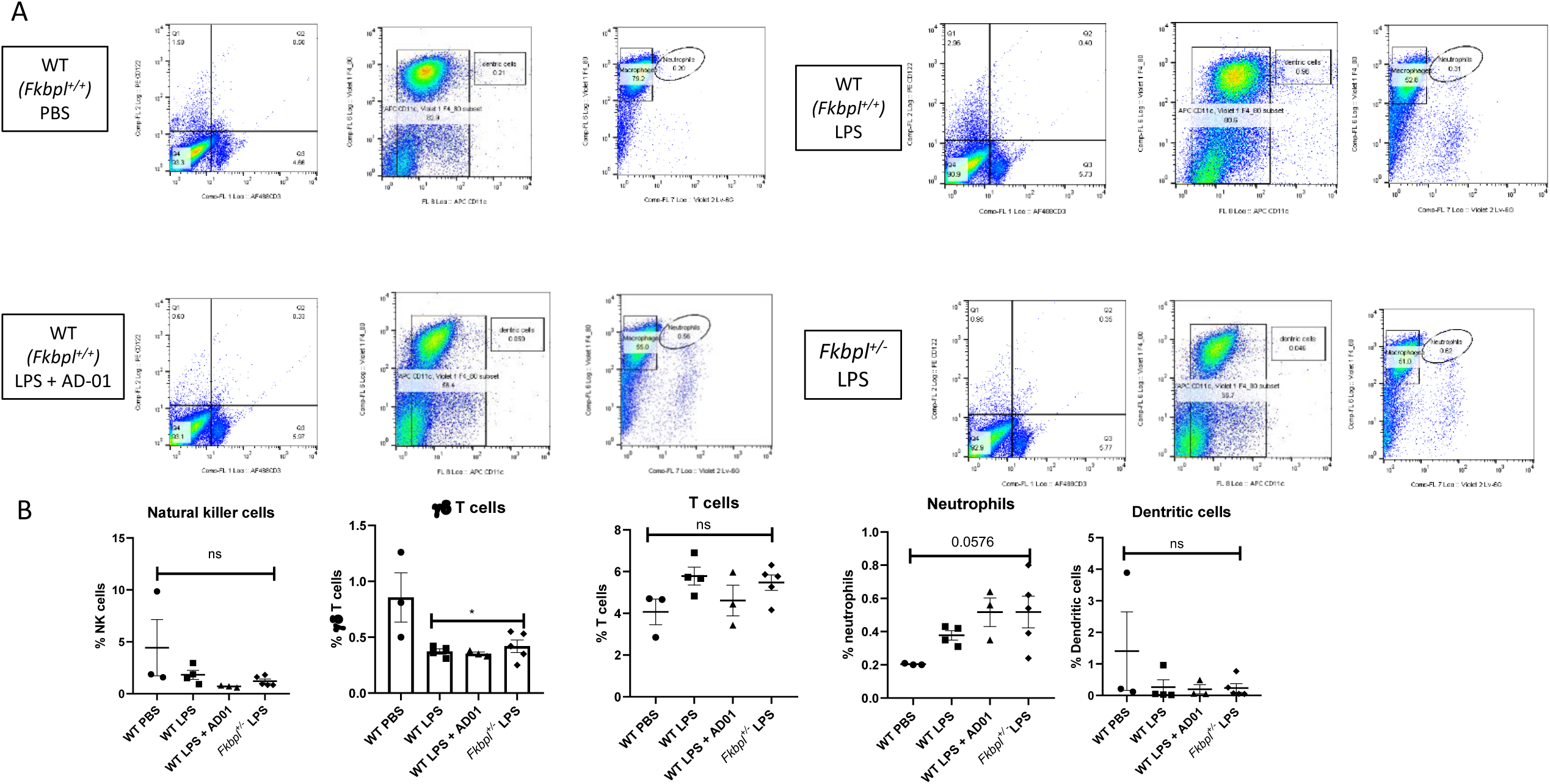
**A)** Flow cytometry gating strategy of immune cell analysis from peritoneal lavage washings collected from wild-type mice treated with PBS, LPS 6 mg/kg and AD-01 3 mg/kg (post LPS) and *Fkbpl*^+/−^ mice treated with LPS 6 mg/ml after 3 h. **(B)** Percentage of natural killer cells, γδ T cells, T cells, neutrophils, dendritic cells in peritoneal lavage washings. Data points are mean ± SEM. n ≥ 3. * p < 0.05, ** p < 0.01, ***p < 0.01 (one way ANOVA).

**Supplementary Figure 3.**
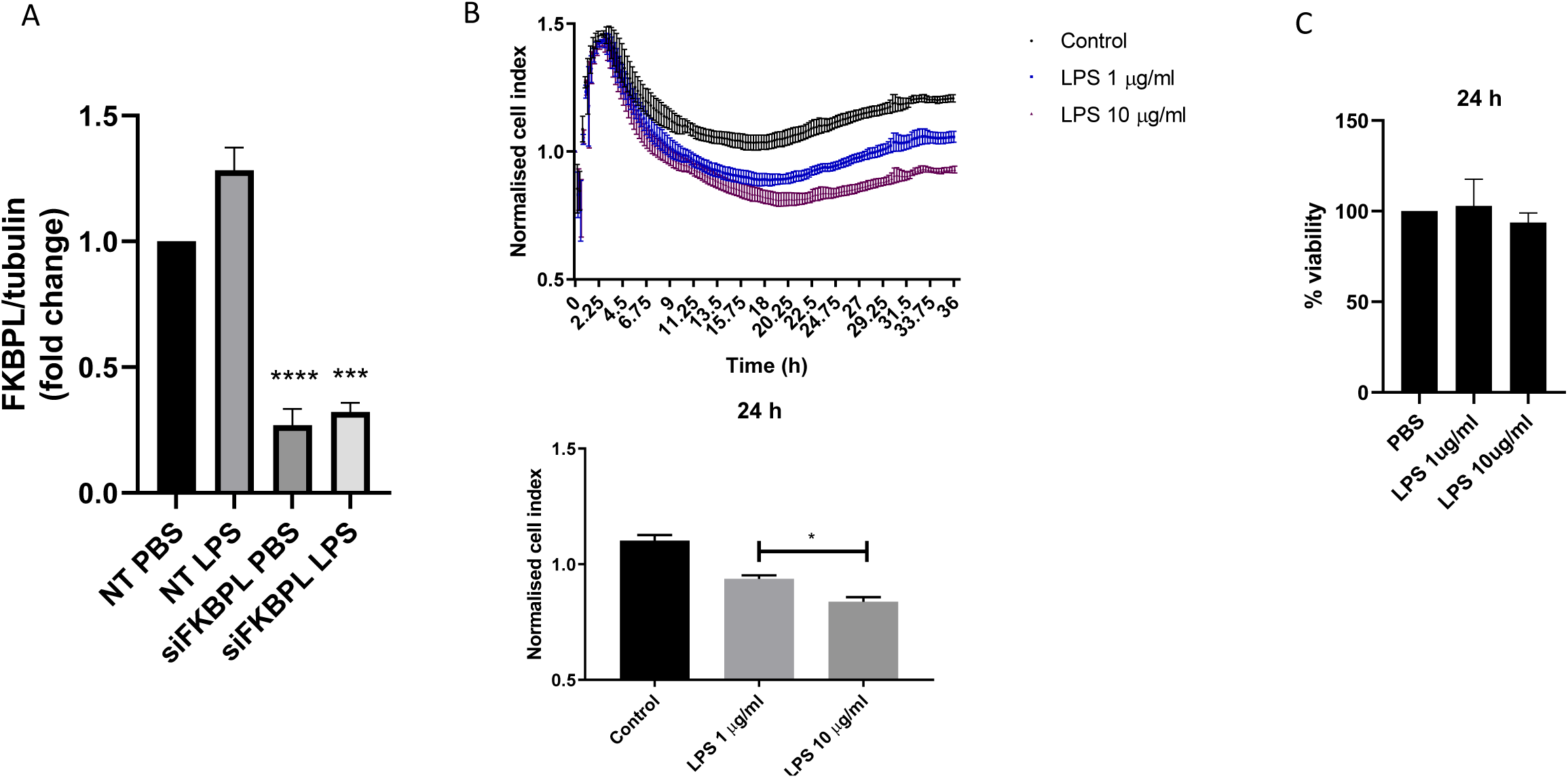
**(A)** FKBPL knockdown in human microvascular endothelial cells (HMEC-1s) using a siFKBPL and non targeted (NT) control in untreated and LPS (10μg/ml) stimulation conditions. Protein FKBPL expression was analyzed by western blot and protein expression was quantified using ImageJ, adjusted to αβ tubulin and normalized to control. **(B)** Cell impedance was measured in HMEC-1s treated with PBS (control) or LPS (1 and 10 μg/ml) using the X-CELLigence system. HMEC-1 cell index for each condition was normalized to PBS control at 24 h. **(C)** Cell viability of HMEC-1s treated with PBS (control) or LPS (1 and 10 μg/ml) for 24 h was measured using MTT assay. Data points are mean ± SEM. n ≥ 3. * p < 0.05, ** p < 0.01, ***p < 0.01 (one way ANOVA).

**Supplementary Figure 4.**
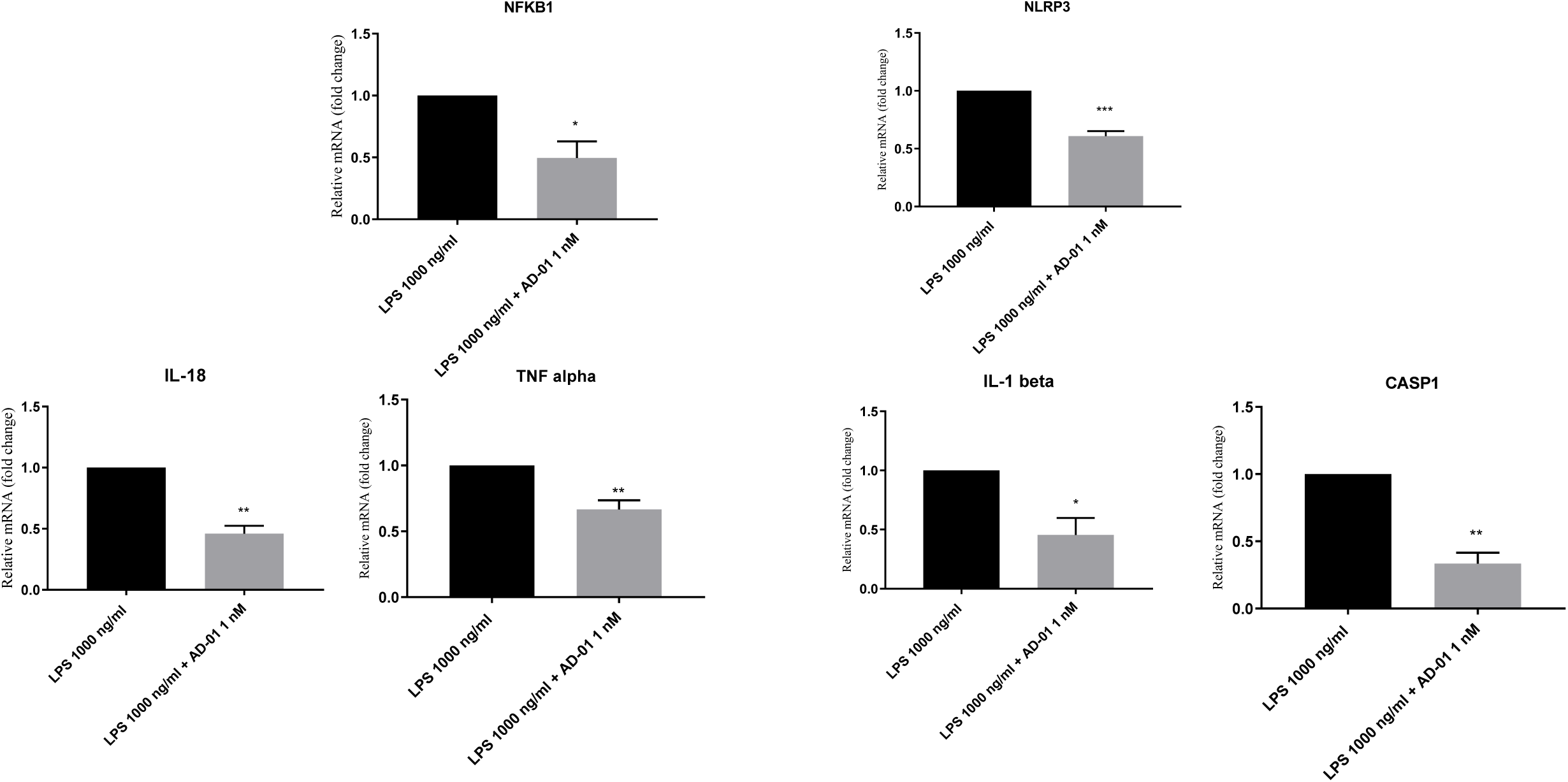
The THP-1 human monocytic cell line was treated with LPS (1000 ng/ml) and AD-01 (1 nM) for 6 h. RNA was extracted and gene expression analysis conducted by qPCR. Data points are mean ± SEM. n ≥ 3. * p < 0.05, ** p < 0.01, ***p < 0.01 (one way ANOVA).

**Supplementary Table 1.** Traits associated with SNPs in FKBP51 and FKBP52

| Protein | Variant | Position | Annotation | P value | Trait |
| --- | --- | --- | --- | --- | --- |
| **FKBP51** | 6:35542476 C / A<br>(rs3800373) | 35542476 | 3' UTR | $6.11 \times 10^{-17}$ | Lymphocyte Count |
| | 6:35553345 A / G<br>(rs73746493) | 35553345 | 3' UTR | $5.01 \times 10^{-16}$ | Lymphocyte Count |
| | 6:35565422 G / A<br>(rs7753746) | 35565422 | intron | $3.49 \times 10^{-18}$ | Eosonophil % |
| | 6:35569281 A / G<br>(rs4713899) | 35569281 | intron | $1.61 \times 10^{-18}$ | Eosonophil % |
| | 6:35589070 C / A<br>(rs9380524) | 35589070 | intron | $8.61 \times 10^{-8}$ | Monocyte % |
| | 6:35607571 T / C<br>(rs1360780) | 35607571 | intron | $1.13 \times 10^{-18}$ | Lymphocyte count |
| | 6:35614328 T / C<br>(rs59595954) | 35614328 | intron | $3.85 \times 10^{-16}$ | Standing Height |
| | 6:35633456 A / G<br>(rs13192954) | 35633456 | intron | $1.01 \times 10^{-11}$ | Trunk fat free mass |
| | 6:35653437 G / A<br>(rs10947563) | 35653437 | intron | $1.21 \times 10^{-9}$ | Non Albumin Protein |
| | 6:35659847 C / A<br>(rs62402145) | 35659847 | intron | $1.77 \times 10^{-10}$ | Trunk free mass |
| | 6:35669983 A / G<br>(rs4713916) | 35669983 | intron | $2.16 \times 10^{-9}$ | Monocyte % |
| | 6:35671651 T / C<br>(rs10456432) | 35671651 | intron | $5.37 \times 10^{-16}$ | FEV1 measure |
| | 6:35672330 G / C<br>(rs9394312) | 35672330 | intron | $2.17 \times 10^{-13}$ | FEV1 measure |
| | 6:35675696 A / G<br>(rs9380529) | 35675696 | intron | $1.86 \times 10^{-13}$ | FEV1 measure |
| | 6:35692928 C / T<br>(rs12203716) | 35692928 | intron | $2.80 \times 10^{-11}$ | Standing Height |
| | 6:35694245 C / T<br>(rs76917354) | 35694245 | intron | $2.48 \times 10^{-12}$ | Standing Height |
| **FKBP52** | 12:2907978 G / T<br>(rs200798844) | 2907978 | missense | $4.92 \times 10^{-13}$ | Velo cardio function |
| | 12:2908330 C / A<br>(rs56196860) | 2908330 | missense | $1.20 \times 10^{-144}$ | Testosterone adjusted |

**Supplementary Table 2.** Gene expression changes predicted from SNP rs28732176

| Gencode Id | Gene Symbol | P-Value | NES | Tissue |
| --- | --- | --- | --- | --- |
| ENSG00000204308.7 | RNF5 | 1.70E-21 | -0.29 | Muscle - Skeletal |
| ENSG00000248290.1 | TNXA | 5.30E-17 | 0.98 | Testis |
| ENSG00000248290.1 | TNXA | 5.60E-17 | 0.72 | Nerve - Tibial |
| ENSG00000248290.1 | TNXA | 4.90E-16 | 0.68 | Adipose - Visceral (Omentum) |
| ENSG00000248290.1 | TNXA | 1.20E-14 | 0.58 | Esophagus - Muscularis |
| ENSG00000248290.1 | TNXA | 2.20E-14 | 1 | Adrenal Gland |
| ENSG00000204315.3 | FKBPL | 1.30E-13 | -0.24 | Skin - Sun Exposed (Lower leg) |
| ENSG00000204315.3 | FKBPL | 1.80E-12 | -0.32 | Skin - Not Sun Exposed (Suprapubic) |
| ENSG00000248290.1 | TNXA | 2.00E-12 | 0.61 | Colon - Transverse |
| ENSG00000204315.3 | FKBPL | 3.00E-12 | 0.33 | Whole Blood |
| ENSG00000204366.3 | ZBTB12 | 4.60E-12 | 0.49 | Whole Blood |
| ENSG00000234745.10 | HLA-B | 5.10E-12 | 0.27 | Whole Blood |
| ENSG00000204516.9 | MICB | 6.80E-12 | -0.37 | Thyroid |
| ENSG00000250535.1 | STK19B | 2.50E-11 | 0.59 | Skin - Sun Exposed (Lower leg) |
| ENSG00000250535.1 | STK19B | 2.60E-11 | 0.51 | Thyroid |
| ENSG00000248290.1 | TNXA | 7.90E-11 | 0.45 | Thyroid |
| ENSG00000250535.1 | STK19B | 1.00E-10 | 0.61 | Adipose - Visceral (Omentum) |
| ENSG00000225851.1 | HLA-S | 1.90E-10 | 0.55 | Whole Blood |
| ENSG00000204516.9 | MICB | 2.20E-10 | -0.43 | Skin - Sun Exposed (Lower leg) |
| ENSG00000250535.1 | STK19B | 2.60E-10 | 0.61 | Nerve - Tibial |
| ENSG00000229391.7 | HLA-DRB6 | 4.90E-10 | 0.54 | Adipose - Subcutaneous |
| ENSG00000250535.1 | STK19B | 1.20E-09 | 0.5 | Whole Blood |
| ENSG00000250535.1 | STK19B | 2.00E-09 | 0.58 | Esophagus - Mucosa |
| ENSG00000204308.7 | RNF5 | 3.90E-09 | -0.23 | Heart - Left Ventricle |
| ENSG00000204301.6 | NOTCH4 | 5.50E-09 | -0.21 | Adipose - Subcutaneous |
| ENSG00000229391.7 | HLA-DRB6 | 7.30E-09 | 0.51 | Nerve - Tibial |
| ENSG00000204314.10 | PRRT1 | 1.30E-08 | 0.24 | Cells - Cultured fibroblasts |
| ENSG00000248290.1 | TNXA | 2.30E-08 | 0.74 | Pituitary |
| ENSG00000204516.9 | MICB | 2.50E-08 | -0.27 | Whole Blood |
| ENSG00000229391.7 | HLA-DRB6 | 3.40E-08 | 0.45 | Muscle - Skeletal |
| ENSG00000232629.8 | HLA-DQB2 | 5.30E-08 | 0.48 | Adipose - Subcutaneous |
| ENSG00000204308.7 | RNF5 | 7.40E-08 | -0.22 | Colon - Transverse |
| ENSG00000248290.1 | TNXA | 7.60E-08 | 0.55 | Stomach |
| ENSG00000250535.1 | STK19B | 7.70E-08 | 0.45 | Adipose - Subcutaneous |
| ENSG00000229391.7 | HLA-DRB6 | 1.10E-07 | 0.38 | Whole Blood |
| ENSG00000250535.1 | STK19B | 1.40E-07 | 0.52 | Breast - Mammary Tissue |
| ENSG00000229391.7 | HLA-DRB6 | 1.40E-07 | 0.45 | Skin - Sun Exposed (Lower leg) |
| ENSG00000248290.1 | TNXA | 1.80E-07 | 0.5 | Breast - Mammary Tissue |
| ENSG00000248290.1 | TNXA | 2.10E-07 | 0.84 | Ovary |
| ENSG00000250535.1 | STK19B | 2.40E-07 | 0.47 | Esophagus - Muscularis |
| ENSG00000204516.9 | MICB | 3.20E-07 | -0.39 | Esophagus - Mucosa |
| ENSG00000232629.8 | HLA-DQB2 | 3.70E-07 | 0.42 | Muscle - Skeletal |
| ENSG00000204344.14 | STK19 | 4.50E-07 | -0.19 | Skin - Sun Exposed (Lower leg) |
| ENSG00000225851.1 | HLA-S | 5.20E-07 | 0.44 | Adipose - Subcutaneous |
| ENSG00000204301.6 | NOTCH4 | 6.30E-07 | -0.19 | Skin - Sun Exposed (Lower leg) |
| ENSG00000250535.1 | STK19B | 6.60E-07 | 0.65 | Testis |
| ENSG00000196126.11 | HLA-DRB1 | 8.00E-07 | -0.33 | Pancreas |
| ENSG00000196126.11 | HLA-DRB1 | 8.40E-07 | -0.25 | Lung |
| ENSG00000168477.17 | TNXB | 8.40E-07 | 0.48 | Adrenal Gland |
| ENSG00000229391.7 | HLA-DRB6 | 8.90E-07 | 0.42 | Esophagus - Mucosa |
| ENSG00000196126.11 | HLA-DRB1 | 9.10E-07 | -0.28 | Breast - Mammary Tissue |
| ENSG00000248290.1 | TNXA | 9.50E-07 | 0.7 | Liver |
| ENSG00000204301.6 | NOTCH4 | 9.70E-07 | -0.24 | Lung |
| ENSG00000225851.1 | HLA-S | 0.000001 | 0.42 | Skin - Sun Exposed (Lower leg) |
| ENSG00000204516.9 | MICB | 1.2E-06 | -0.36 | Esophagus - Muscularis |
| ENSG00000204536.13 | CCHCR1 | 1.2E-06 | 0.25 | Nerve - Tibial |
| ENSG00000204308.7 | RNF5 | 1.2E-06 | -0.35 | Adrenal Gland |
| ENSG00000234745.10 | HLA-B | 1.5E-06 | 0.25 | Nerve - Tibial |
| ENSG00000204308.7 | RNF5 | 1.7E-06 | -0.19 | Heart - Atrial Appendage |
| ENSG00000250535.1 | STK19B | 1.8E-06 | 0.78 | Ovary |
| ENSG00000204301.6 | NOTCH4 | 1.9E-06 | -0.27 | Esophagus - Muscularis |
| ENSG00000248290.1 | TNXA | 1.9E-06 | 0.38 | Lung |
| ENSG00000234745.10 | HLA-B | 3.3E-06 | 0.2 | Adipose - Subcutaneous |
| ENSG00000248290.1 | TNXA | 3.3E-06 | 0.45 | Esophagus - Gastroesophageal Junction |
| ENSG00000204308.7 | RNF5 | 3.5E-06 | -0.19 | Thyroid |
| ENSG00000204301.6 | NOTCH4 | 3.7E-06 | -0.24 | Heart - Atrial Appendage |
| ENSG00000234745.10 | HLA-B | 0.000004 | 0.21 | Thyroid |
| ENSG00000204531.17 | POU5F1 | 4.4E-06 | 0.34 | Adipose - Subcutaneous |
| ENSG00000204438.10 | GPANK1 | 4.4E-06 | 0.13 | Whole Blood |
| ENSG00000204438.10 | GPANK1 | 4.8E-06 | 0.2 | Adipose - Subcutaneous |
| ENSG00000250535.1 | STK19B | 5.2E-06 | 0.6 | Esophagus - Gastroesophageal Junction |
| ENSG00000250535.1 | STK19B | 5.3E-06 | 0.5 | Colon - Transverse |
| ENSG00000248290.1 | TNXA | 5.9E-06 | 0.55 | Colon - Sigmoid |
| ENSG00000232629.8 | HLA-DQB2 | 0.000006 | 0.48 | Adipose - Visceral (Omentum) |
| ENSG00000250535.1 | STK19B | 6.1E-06 | 0.51 | Pancreas |
| ENSG00000204516.9 | MICB | 6.2E-06 | -0.31 | Adipose - Subcutaneous |
| ENSG00000204538.3 | PSORS1C2 | 7.1E-06 | 0.42 | Nerve - Tibial |
| ENSG00000204308.7 | RNF5 | 7.1E-06 | -0.33 | Pancreas |
| ENSG00000196126.11 | HLA-DRB1 | 7.8E-06 | -0.24 | Artery - Aorta |
| ENSG00000234745.10 | HLA-B | 7.8E-06 | 0.31 | Esophagus - Gastroesophageal Junction |
| ENSG00000204538.3 | PSORS1C2 | 7.9E-06 | 0.44 | Adipose - Visceral (Omentum) |
| ENSG00000229391.7 | HLA-DRB6 | 8.8E-06 | 0.36 | Thyroid |
| ENSG00000204538.3 | PSORS1C2 | 9.6E-06 | 0.41 | Artery - Tibial |
| ENSG00000196126.11 | HLA-DRB1 | 9.7E-06 | -0.22 | Skin - Sun Exposed (Lower leg) |
| ENSG00000204301.6 | NOTCH4 | 0.000011 | -0.24 | Nerve - Tibial |
| ENSG00000234745.10 | HLA-B | 0.000011 | 0.19 | Heart - Left Ventricle |
| ENSG00000204385.10 | SLC44A4 | 0.000011 | 0.23 | Pituitary |
| ENSG00000204256.12 | BRD2 | 0.000013 | -0.22 | Skin - Not Sun Exposed (Suprapubic) |
| ENSG00000248290.1 | TNXA | 0.000016 | 0.8 | Brain - Hypothalamus |
| ENSG00000225851.1 | HLA-S | 0.000016 | 0.83 | Uterus |
| ENSG00000229391.7 | HLA-DRB6 | 0.000016 | 0.43 | Esophagus - Muscularis |
| ENSG00000234745.10 | HLA-B | 0.000017 | 0.25 | Artery - Aorta |
| ENSG00000229391.7 | HLA-DRB6 | 0.000017 | 0.62 | Prostate |
| ENSG00000204301.6 | NOTCH4 | 0.000017 | -0.21 | Skin - Not Sun Exposed (Suprapubic) |
| ENSG00000221988.12 | PPT2 | 0.000017 | 0.19 | Cells - Cultured fibroblasts |
| ENSG00000250535.1 | STK19B | 0.000018 | 0.38 | Artery - Tibial |
| ENSG00000229391.7 | HLA-DRB6 | 0.000019 | 0.44 | Skin - Not Sun Exposed (Suprapubic) |
| ENSG00000250535.1 | STK19B | 0.000019 | 0.47 | Artery - Aorta |
| ENSG00000232629.8 | HLA-DQB2 | 0.00002 | 0.4 | Nerve - Tibial |
| ENSG00000229391.7 | HLA-DRB6 | 0.000022 | 0.45 | Adipose - Visceral (Omentum) |
| ENSG00000204516.9 | MICB | 0.000023 | -0.28 | Cells - Cultured fibroblasts |
| ENSG00000204301.6 | NOTCH4 | 0.000026 | -0.22 | Adipose - Visceral (Omentum) |
| ENSG00000204538.3 | PSORS1C2 | 0.000028 | 0.53 | Esophagus - Gastroesophageal Junction |
| ENSG00000204538.3 | PSORS1C2 | 0.000031 | 0.34 | Adipose - Subcutaneous |
| ENSG00000204421.2 | LY6G6C | 0.000033 | -0.28 | Esophagus - Mucosa |
| ENSG00000237541.3 | HLA-DQA2 | 0.000036 | 0.34 | Whole Blood |
| ENSG00000272221.1 | XXbac-BPG181B23.7 | 0.000042 | -0.35 | Skin - Sun Exposed (Lower leg) |
| ENSG00000204516.9 | MICB | 0.000042 | -0.34 | Nerve - Tibial |
| ENSG00000196126.11 | HLA-DRB1 | 0.000045 | -0.39 | Small Intestine - Terminal Ileum |
| ENSG00000232629.8 | HLA-DQB2 | 0.000047 | 0.54 | Colon - Sigmoid |
| ENSG00000237541.3 | HLA-DQA2 | 0.000051 | 0.4 | Nerve - Tibial |
| ENSG00000225851.1 | HLA-S | 0.000051 | 0.43 | Adipose - Visceral (Omentum) |
| ENSG00000241106.6 | HLA-DOB | 0.000056 | -0.32 | Artery - Tibial |
| ENSG00000196126.11 | HLA-DRB1 | 0.000064 | -0.21 | Thyroid |
| ENSG00000204516.9 | MICB | 0.000066 | -0.34 | Heart - Atrial Appendage |
| ENSG00000248290.1 | TNXA | 0.000068 | 0.64 | Small Intestine - Terminal Ileum |
| ENSG00000213654.9 | GPSM3 | 0.00007 | 0.43 | Brain - Cerebellum |
| ENSG00000204516.9 | MICB | 0.000076 | -0.26 | Breast - Mammary Tissue |
| ENSG00000204531.17 | POU5F1 | 0.000076 | 0.36 | Nerve - Tibial |
| ENSG00000250535.1 | STK19B | 0.000078 | 0.35 | Cells - Cultured fibroblasts |
| ENSG00000248290.1 | TNXA | 0.000081 | 0.64 | Spleen |
| ENSG00000196126.11 | HLA-DRB1 | 0.000083 | -0.19 | Esophagus - Mucosa |
| ENSG00000204301.6 | NOTCH4 | 0.000087 | -0.16 | Artery - Tibial |
| ENSG00000232629.8 | HLA-DQB2 | 0.000088 | 0.31 | Thyroid |
| ENSG00000204301.6 | NOTCH4 | 0.00009 | -0.18 | Heart - Left Ventricle |
| ENSG00000196126.11 | HLA-DRB1 | 0.000094 | -0.23 | Skin - Not Sun Exposed (Suprapubic) |
| ENSG00000204538.3 | PSORS1C2 | 0.000097 | 0.38 | Lung |
| ENSG00000204344.14 | STK19 | 0.0001 | -0.22 | Nerve - Tibial |
| ENSG00000250535.1 | STK19B | 0.00011 | 0.78 | Brain - Cerebellar Hemisphere |
| ENSG00000204516.9 | MICB | 0.00011 | -0.23 | Lung |
| ENSG00000204516.9 | MICB | 0.00011 | -0.3 | Artery - Tibial |
| ENSG00000204308.7 | RNF5 | 0.00012 | -0.15 | Esophagus - Muscularis |
| ENSG00000232629.8 | HLA-DQB2 | 0.00012 | 0.4 | Esophagus - Muscularis |
| ENSG00000204310.12 | AGPAT1 | 0.00012 | -0.13 | Muscle - Skeletal |
| ENSG00000204301.6 | NOTCH4 | 0.00012 | -0.3 | Testis |
| ENSG00000248290.1 | TNXA | 0.00012 | 0.31 | Esophagus - Mucosa |
| ENSG00000229391.7 | HLA-DRB6 | 0.00013 | 0.4 | Testis |
| ENSG00000226979.8 | LTA | 0.00013 | 0.28 | Testis |
| ENSG00000204301.6 | NOTCH4 | 0.00014 | -0.22 | Esophagus - Mucosa |
| ENSG00000204410.14 | MSH5 | 0.00014 | 0.14 | Esophagus - Mucosa |
| ENSG00000237541.3 | HLA-DQA2 | 0.00015 | 0.35 | Adipose - Subcutaneous |
| ENSG00000198563.13 | DDX39B | 0.00016 | 0.14 | Nerve - Tibial |
| ENSG00000204536.13 | CCHCR1 | 0.00017 | 0.17 | Whole Blood |
| ENSG00000204516.9 | MICB | 0.00018 | -0.25 | Adipose - Visceral (Omentum) |
| ENSG00000204538.3 | PSORS1C2 | 0.00018 | 0.31 | Cells - Cultured fibroblasts |
| ENSG00000232629.8 | HLA-DQB2 | 0.0002 | 0.27 | Whole Blood |
| ENSG00000204392.10 | LSM2 | 0.00021 | -0.18 | Esophagus - Muscularis |
| ENSG00000204410.14 | MSH5 | 0.00021 | 0.12 | Skin - Sun Exposed (Lower leg) |
| ENSG00000196126.11 | HLA-DRB1 | 0.00022 | -0.18 | Muscle - Skeletal |
| ENSG00000204338.8 | CYP21A1P | 0.00023 | 0.29 | Adipose - Subcutaneous |
| ENSG00000204516.9 | MICB | 0.00024 | -0.32 | Skin - Not Sun Exposed (Suprapubic) |
| ENSG00000204338.8 | CYP21A1P | 0.0003 | 0.3 | Skin - Sun Exposed (Lower leg) |
| ENSG00000225851.1 | HLA-S | 0.00034 | 0.3 | Thyroid |
| ENSG00000204525.16 | HLA-C | 0.00039 | 0.19 | Adipose - Subcutaneous |
| ENSG00000204516.9 | MICB | 0.00043 | -0.25 | Muscle - Skeletal |

*NES – Normalised Effect Size*

**Supplementary Table 3.** Gene expression changes predicted from SNP rs35580488

| **Gencode Id** | **Gene Symbol** | **P-Value** | **NES** | **Tissue** |
| --- | --- | --- | --- | --- |
| ENSG00000204351.11 | SKIV2L | 1.90E-07 | 0.82 | Esophagus - Muscularis |
| ENSG00000204396.10 | VWA7 | 3.40E-07 | -0.64 | Heart - Atrial Appendage |
| ENSG00000204351.11 | SKIV2L | 1.2E-06 | 0.72 | Skin - Not Sun Exposed (Suprapubic) |
| ENSG00000224389.8 | C4B | 1.2E-06 | -0.96 | Nerve - Tibial |
| ENSG00000224389.8 | C4B | 2.4E-06 | -2.8 | Brain - Amygdala |
| ENSG00000224389.8 | C4B | 5.8E-06 | -2.6 | Brain - Anterior cingulate cortex (BA24) |
| ENSG00000204351.11 | SKIV2L | 0.000014 | 0.79 | Artery - Tibial |
| ENSG00000248290.1 | TNXA | 0.000031 | 0.8 | Adipose - Subcutaneous |
| ENSG00000204351.11 | SKIV2L | 0.000035 | 1.9 | Small Intestine - Terminal Ileum |
| ENSG00000204351.11 | SKIV2L | 0.00004 | 0.56 | Thyroid |
| ENSG00000204351.11 | SKIV2L | 0.000046 | 0.99 | Adrenal Gland |
| ENSG00000204351.11 | SKIV2L | 0.000073 | 0.34 | Whole Blood |
| ENSG00000224389.8 | C4B | 0.000079 | -0.67 | Thyroid |
| ENSG00000204351.11 | SKIV2L | 0.000085 | 0.46 | Cells - Cultured fibroblasts |
| ENSG00000204351.11 | SKIV2L | 0.000099 | 0.58 | Lung |
| ENSG00000204351.11 | SKIV2L | 0.0001 | 0.81 | Colon - Sigmoid |
| ENSG00000204315.3 | FKBPL | 0.00014 | -0.76 | Adipose - Subcutaneous |
| ENSG00000248290.1 | TNXA | 0.00019 | 0.98 | Artery - Tibial |
| ENSG00000204396.10 | VWA7 | 0.0003 | -0.43 | Muscle - Skeletal |

*NES – Normalised Effect Size*

**Supplementary Table 4.**
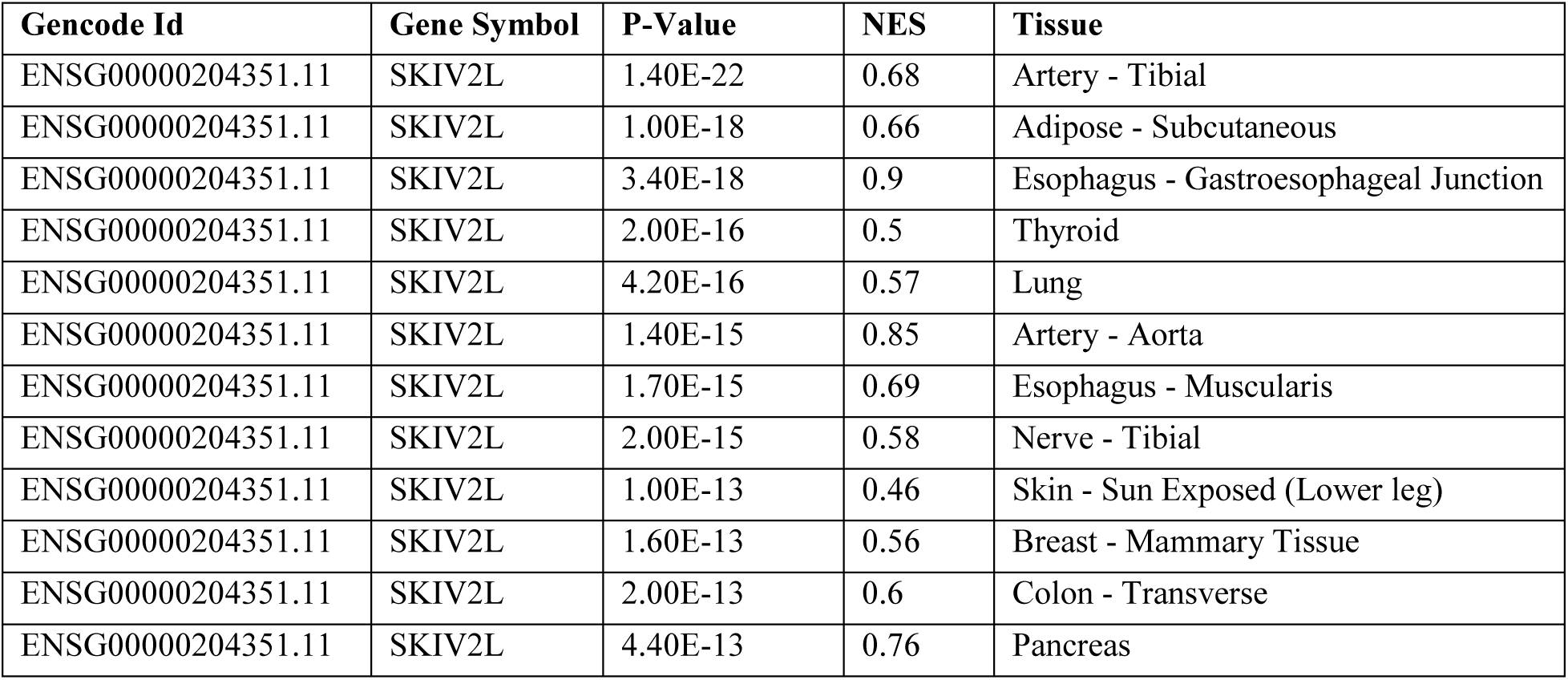
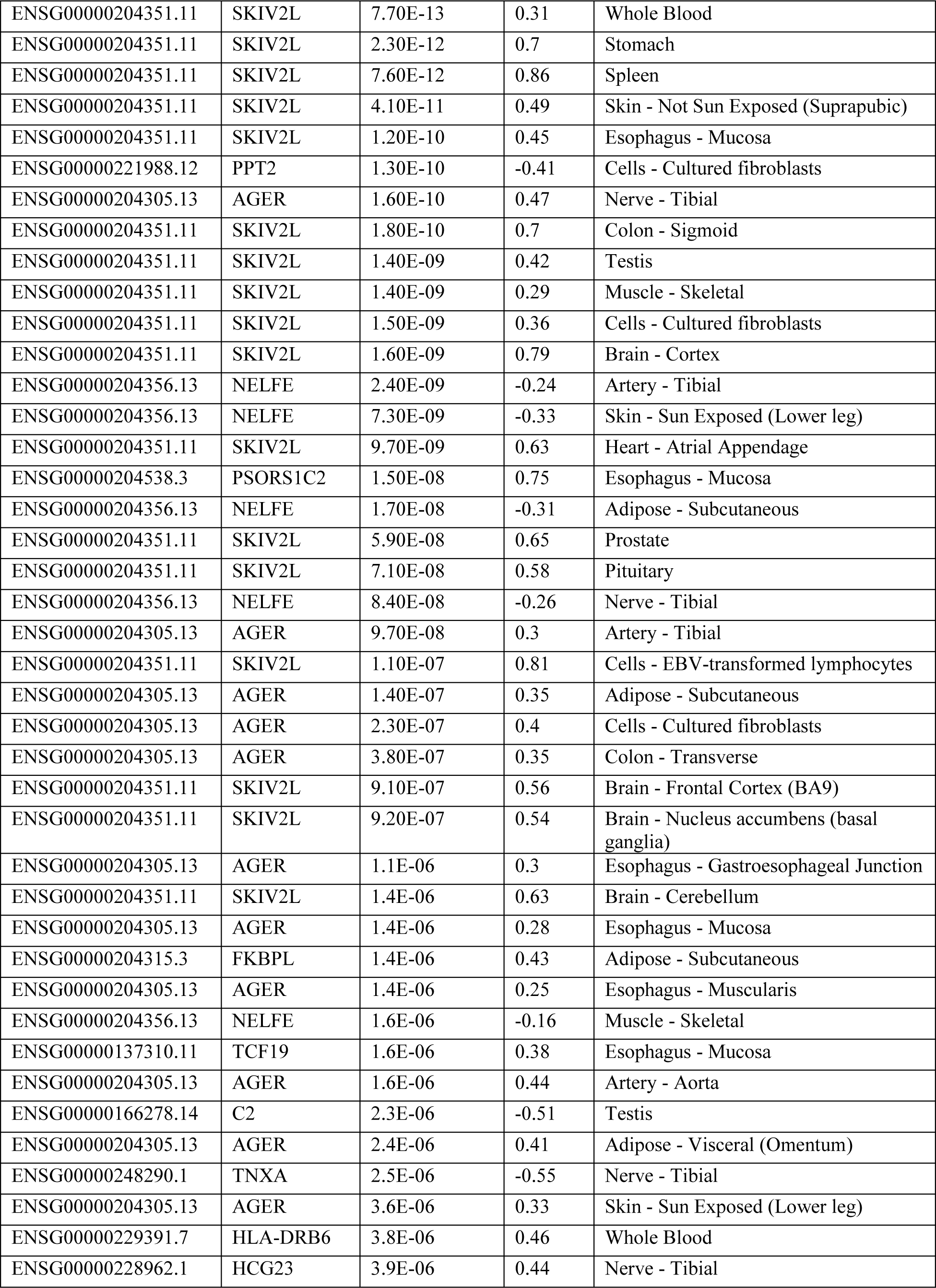
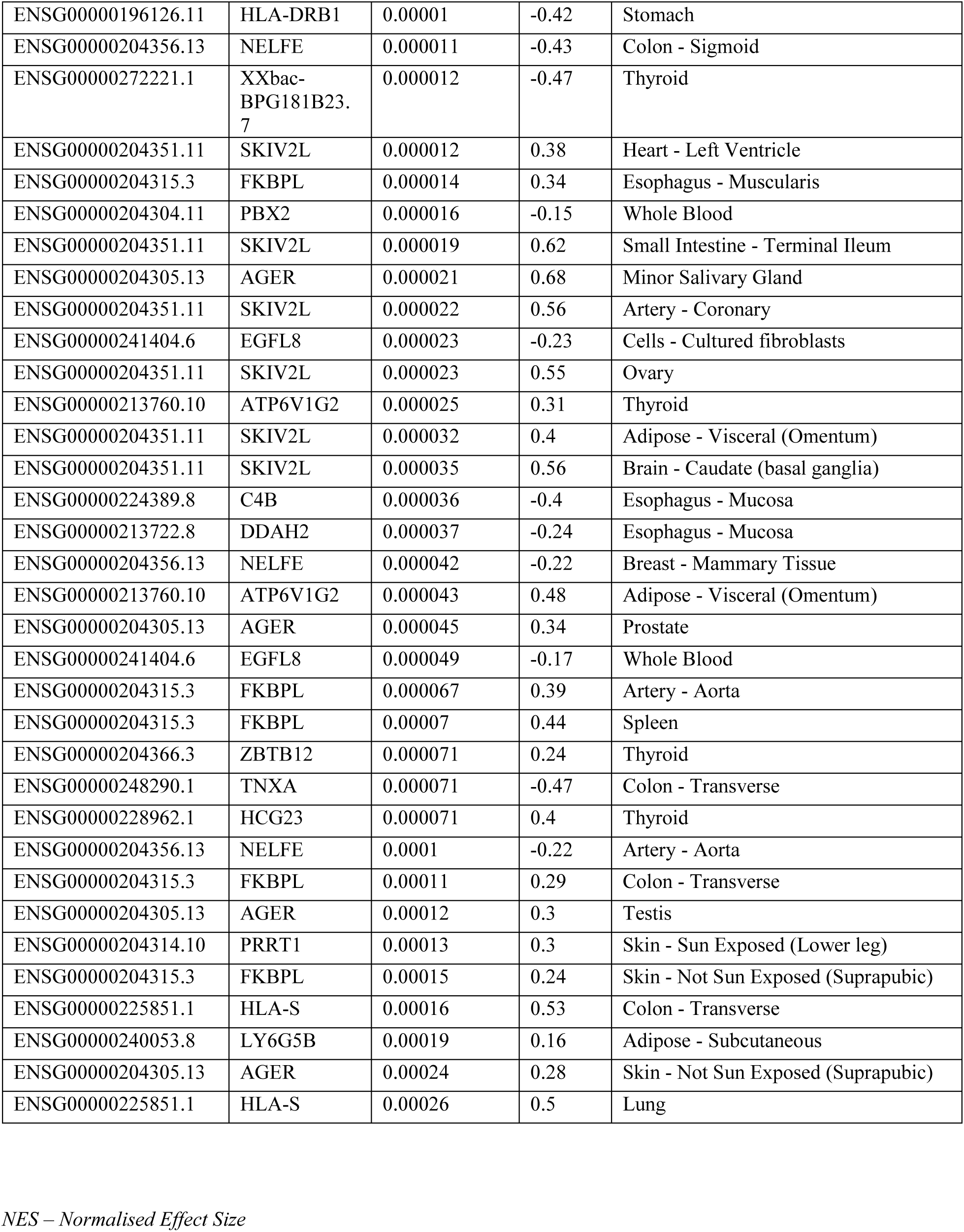
Gene expression changes predicted from SNP rs204892

